# Rejuvenation of Senescent Cells, in vitro and in vivo, by Low-frequency Ultrasound without Senolysis

**DOI:** 10.1101/2022.12.08.519320

**Authors:** Sanjay Kureel, Rosario Maroto, Simon Powell, Maisha Aniqua, Felix Margadant, Brandon Blair, Blake B. Rasmussen, Michael Sheetz

## Abstract

The presence of an appreciable number of senescent cells causes age-related pathologies as their removal by genetic or pharmacological means, as well as possibly by exercise, improves outcome in animal models. An alternative to depleting such cells would be to rejuvenate them to promote their return to a replicative state. Means to do so have not been explored, but here we report that treatment of non-growing senescent cells with low-frequency ultrasound (LFU) rejuvenates the cells. Notably, we find 15 characteristics of senescent cells that are reversed by LFU, including decreased cell and organelle motility. Mechanistically, LFU causes Ca^2+^ entry that precedes dramatic increases in autophagy and an inhibition of mTORC1 signaling and the movement of Sirtuin1 from the nucleus to cytoplasm. Also, there is inhibition of SASP secretion, as well as β-galactosidase expression, telomere length is increased, while nuclear 5mc, H3K9me3, γH2AX, nuclear p53, ROS and mitoROS levels are all restored to normal levels. Repeated LFU treatments enables expansion of primary cells and stem cells beyond normal replicative limits without altering their phenotype. The rejuvenation process is enhanced by co-treatment with rapamycin or Rho kinase inhibition but is inhibited by blocking Sirtuin1 or Piezo1 activity. We further optimized the LFU treatment parameters to increase mouse lifespan and healthspan. These results suggest that mechanically-induced pressure waves alone can reverse senescence and aging effects at the cellular and organismal level, providing a non-pharmacological way to treat the effects of aging.

**One-Sentence Summary:** Low frequency ultrasound is sufficient to rejuvenate senescent cells by activating autophagy in aged mice to improve healthspan.

## Introduction

Cell senescence is one of the hallmarks of the aging process that is purportedly defined by a permanent block to cell growth.^1,2^ It was first noted in 1961 by Hayflick and Moorhead that primary cells in culture stopped growing after a certain number of divisions; that is, they became senescent.^3^ Transplantation of senescent cells into young mice causes physical deterioration and age-related pathologies ^4^, whereas depletion of these cells from aged mice genetically or by the use of senolytics slows the development of age-related pathologies and enhances lifespan.^5–13^ These results indicate that cellular senescence is one of the drivers of the aging process and removing senescent cells is critical for improving performance in aged organisms. Because senescence is believed to be a state of permanent cell cycle arrest in which cells are metabolically active but have stopped dividing ^14^, the selective lysis of senescent cells is considered the best way to remove their negative effects.^15–19^

While cell senescence is typically considered detrimental, it should be noted that the senescence process has general physiological significance as it prevents the propagation of damaged cells, suppresses tumor progression, helps in early development ^20^, wound healing ^21^ and in the tissue repair processes.^22^ Despite the beneficial roles of senescent cells, they are linked to many age-associated maladies ^23–25^, including in the lung, adipose tissue, aorta, pancreas and osteoarthritic joints.^26–28^ Thus, there are potentially many benefits from decreasing the fraction of senescent cells in various tissues during aging. Senescent cells secrete pro-inflammatory molecules, growth factors, chemokines, extracellular matrices, proteases and cytokines, collectively known as the senescence-associated secretary phenotype (SASP).^29,30^ An increase in the level of SASP catalyzes many age-related problems.^31,32^ Therefore, targeted elimination of senescent cells by senolytics can potentially improve age-associated pathologies, including osteoarthritis^9^, diabetes^33^, osteoporosis, neurodegenerative disease^34^ and overall lifespan^5^. Senolytics have received considerable attention as a way to diminish specific effects of aging in tissues targeted by them ^35,36^, and they have entered clinical trials. However, there is currently no approved senolytic-based treatment for humans.^6,37,38^

Autophagy inhibition and mitochondrial dysfunction are hallmarks of aging and cellular senescence.^1,39^ Dynamic changes in mitochondrial fusion and fission are essential for healthy mitochondrial function and increased fusion contributes to cell senescence.^40^ There is evidence of decreased senescence with exercise ^41^ that may be related to unknown processes that increase mitochondrial fission.^40,42^ Although changes in lysosome and mitochondrial functions may be related to each other, there is significant evidence that inhibition of lysosomal autophagy is a driver for aging.^43–45^

Mutations in the genome can increase the lifespan of worms and flies and they are largely linked to metabolism and proteostasis pathways that inhibit the onset of senescence but do not reverse senescence at the cellular level.^46^ Strategies to prolong lifespan, including the use of small molecules (*e*.*g*., rapamycin and metformin) and caloric restriction, often can activate autophagy.

An alternative to senolysis is to block the transition to the senescent state, but such an intervention is required relatively early in the aging process and is not applicable to aged individuals with senescent cells. Even so, physical exercise is widely accepted as an early intervention that slows natural aging ^47–49^ and has even been considered as a senolytic.^17^ Although the effects of exercise could be purely physical, there is evidence that exercise affects the brain and other organs through increased secretion of myokines, such as irisin.^50^

Ultrasound produces short duration pressure waves in cells that create mechanical stresses. These conditions are safe for normal tissues and do not adversely influence functions of normal cells.^51^ Previously, we found that ultrasound could activate apoptosis selectively in tumor cells; and it was logical to test if it had similar effects on senescent cells. When we tested ultrasound on senescent cells, surprisingly, low-frequency ultrasound (LFU) treatment restored the growth of chemically induced and replicative senescent cells. Mechanistically, we found that LFU activated autophagy, organelle and cell motility, while it inhibited mTORC1, SASP secretion, β-galactosidase expression, and it decreased cell size and mitochondrial length. Notably, we also found that normal cells treated with LFU secreted factor(s) that activated the growth of senescent cells. Thus, purely mechanical stimuli can selectively rejuvenate senescent cells.

## RESULTS

### Development and optimization of LFU

As previous studies showed that LFU caused apoptosis of cancer cells ^52,53^, we treated senescent Vero cells with an LFU apparatus (Figure S1A, B) to determine if they behaved similarly. Surprisingly, the senescent cells grew and increased in number after LFU treatment rather than undergoing apoptosis (Fig. S1C-F). The LFU bath had a peak power at the center and the cells were positioned 7-10 cm above the ultrasound transducer to put them in the far field of the ultrasound (Figure S1A, B). After testing different ultrasound frequencies and power levels for their effects on the growth of the senescent Vero cells, we found that growth was greater at an LFU of 33 kHz than at 39 kHz and peaked at a power level of 4 Pa as measured by a calibrated hydrophone (Figure S1C, D). Similarly, we tested the effects of different LFU durations and duty cycles (on/off times) on the growth rate of the senescent cells (Figure S1E, F). In this manner we determined that an LFU of 32.2 kHz at 4 Pa for 30 mins with a duty cycle of 1.5s on and 1.5s off had the optimal parameters for inducing the growth of the senescent cells, and thus these values were used for all subsequent cell and mouse experiments.

To test the generalizability of LFU treatments for reversing cell senescence, we induced cell senescence by treating the cells with either doxorubicin (Dox), hydrogen peroxide (H_2_O_2_), sodium butyrate (SB) or bleomycin sulfate (BS) for 36-48 hours followed by 4 days of incubation in medium (*59*). Senescence was characterized by a block of cell growth (statistically, not different from zero growth), greater β-galactosidase activity (>90% of cells were positive), a larger cell size and production of a SASP that inhibited the growth of normal cells. ^28,29^ All four criteria were met with each type of drug-mediated induction of cell senescence cell (Fig. S2A-G). Thus, by all criteria, each of the 4 drugs were able to induce senescence in Vero cells.

### LFU rejuvenates senescent cells without senolysis

To determine if LFU treatment could reverse all types of the senescent cells, we treated Vero cells induced to be senescent by different stresses (H_2_O_2_, BS, Dox or SB) with LFU at the optimal power levels (Figure S3A). After LFU exposure for thirty minutes, the senescent cells induced by all four stressors significantly grew (Figures S3B and S4A-C) and decreased in area (Figures S3C and S4D-F). After the first passage, the growth rate of SB-treated cells actually exceeded that of the parental, non-senescent line (Figure S3B). We also found that while LFU caused the greatest increase in growth rate with SB treated cells (Fig. S3B), LFU also caused significant increases in the number of cells treated with H_2_O_2_, BS, and Dox (Fig. S4A-C). In the first two days, large cells divided, and the distribution of cell sizes shifted to smaller sizes (Figures S3C and S4D-F), indicating that cells became smaller with growth, reaching a normal cell size by the third passage (Figure S3C). When the senescent cells were analyzed for apoptosis by Annexin 5 binding after LFU treatment, there was only background levels of staining, indicating that LFU did not have a senolytic effect (Fig. S4G and H). After a single 30 minute exposure to LFU, the cells grew until at least passage 3 without a significant loss in their growth rate as determined by cell counting (Figure S3B). LFU treated cells had a reduced level of cell cycle inhibitor p21 (Figure S3D and S3E). Thus, it appears that LFU treatment activated growth of senescent cells without senolysis.

### Senescent cells are rejuvenated for growth by LFU

It was possible that the drug-treated cells were quiescent with only a fraction being partially senescent. However, quiescent cells are characterized by an absence of β-galactosidase activity, and we found that in our conditions over 95% of cells stained for β-galactosidase activity after a 4-day incubation in media (Figure S2F,G).^54^ Thus, our data strongly indicates a lack of an appreciable fraction of quiescent cells in these preparations. But to further test if the cells were senescent and if the senescent cells were reactivatable, the cells were analyzed by video microscopy. In particular, BS-treated cells were incubated for 22 days in media after a 2-day treatment with the drug to assure that they were senescent before videotaping them for 24 hours, which enabled the tracking of each cell (see plan Figure 1A). During the 24 hours of taping, there were no mitotic events and limited motility (Figure 1B) but after LFU treatment for 30’, there was significant motility and cell growth (Figure 1C). By following the cells in the videos, it was clear that the senescent cells did not divide before; but after LFU treatment, they increased significantly in number (Figure 1D). The velocity of cell migration of the senescent cells was low as measured by the hourly displacements of the nuclei but after LFU treatment, the velocity of the same cells increased two fold (Figure 1E). Thus, fully senescent cells were activated for growth and motility by LFU.

**Figure 1.**
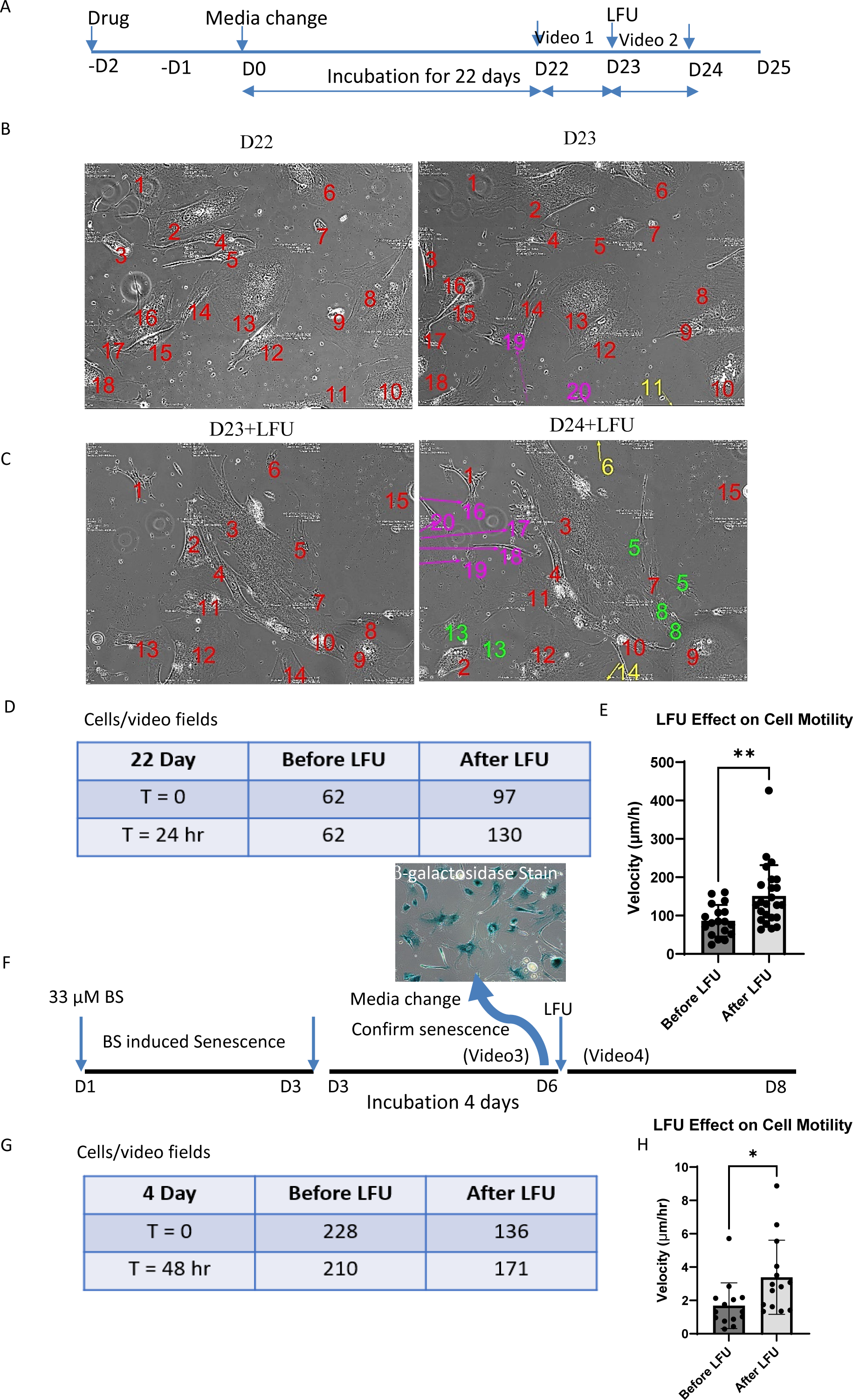
Fully Senescent Cells are Rejuvenated for Growth and Motility by LFU. (A) This diagram shows the treatment plan to produce cells that are fully senescent. Note the timing of the videos 1 and 2 are for 24 hr before and 48 hr after LFU treatment for 30’. (B) Time-lapse video images of a field with 18 Bleomycin sulfate (BS) treated cells incubated for 22 days in vitro, at the beginning and end of a 24 hour observation period (red numbers) where images were taken every 30’ (two cells that entered the field in that period are marked by purple numbers and one that left by yellow). (C) Time-lapse images of a field of 14 cells after a 30’ LFU treatment at the beginning and end of a 24 hour observation period (red numbers). Three of the orginal cells divided (#s 5, 8 and 13) and the daughter cells are noted by green numbers. Five cells entered the field (Purple numbers) and two left the field (yellow numbers). (D) Twenty-eight image fields were tracked for 24 hours before and for 48 hours after LFU treatment and the number of cells in those fields were counted. (E) Cell velocity (determined by displacements of nuclei over each hour for 24 hr.). (F) Diagram of 4 day incubation of BS stressed cells with an image of b-galactosidase staining of the cells after four days of incubation. Note the timing of the videos 3 and 4 are for 48 hr before and after LFU treatment. (G) Twenty-eight image fields were tracked for 48 hr before and after LFU treatment and the number of cells in the fields were counted. (H) The velocity of the cell movements was determined by measuring the displacements of the nuclei over each hour of imaging for 24 hours. Results are shown as mean ± SD, minimally 108 cells were analyzed, n>3 experiments, and significance was determined using two-tailed unpaired t-test. *P<0.05, ** P<0.002.

As a further test of rejuvenation, we followed BS-treated cells that were incubated in media for four days after a 2-day drug treatment (Figure 1F), as such an incubation time period has been suggested to be sufficient to enable such treated cells to move to a senescent state.^55^ In the 4-day incubation protocol, as compared to vehicle-treated cells, all the BS-treated cells expressed β-galactosidase after four days and the time lapse videos showed that there were fewer cells after 48 hours, indicating that some of the drug-treated cells underwent apoptosis (Figure 1G). After LFU, the drug-treated cells grew and there was a similar percentage increase in the cell number as in the 22 day incubated cells. Also, after LFU there was a significantly greater degree of cell motility compared to the before-LFU condition (Figure 1H), suggesting the senescent state was reversed.

### LFU activates mitochondrial dynamics and motility along with lysosomes

To determine if organelle motility was activated as well as nuclear motility, we followed the movements of the mitochondria. The mitochondria of senescent cells were often fused and were longer than those in normal, dividing cells (Figure 2A). LFU treatment caused the fragmentation of the mitochondria and a decrease in the lysosome staining intensity (Figure 2B and C). To determine the velocities of movement, the positions of Mitotracker-labelled mitochondria were recorded every 5-10s. After LFU treatment, the mitochondria were significantly smaller (Figure 2A and C) and the linear velocities were significantly greater (by over 5-fold) than before the cells were exposed to LFU (Figure 2D). In addition, there were many mitochondrial fission and fusion events after LFU (data not shown). For consistency, we only recorded the velocities of mitochondria that did not undergo fission or fusion during the observation period. Lysosomes in the same cells decreased in staining intensity with lysotracker and moved from the center of the cell toward the periphery as the lysosomes potentially were activated for autophagy (Figure 2A and B) giving rise to the hypothesis that LFU caused rejuvenation by activating autophagy through phagosome fusion with lysosomes (Figure 2E). Thus, LFU treatment activated whole-cell, lysosomal and mitochondrial dynamics, indicating that in senescent cells both actin- and microtubule-based motility were promoted after LFU exposure.

**Figure 2.**
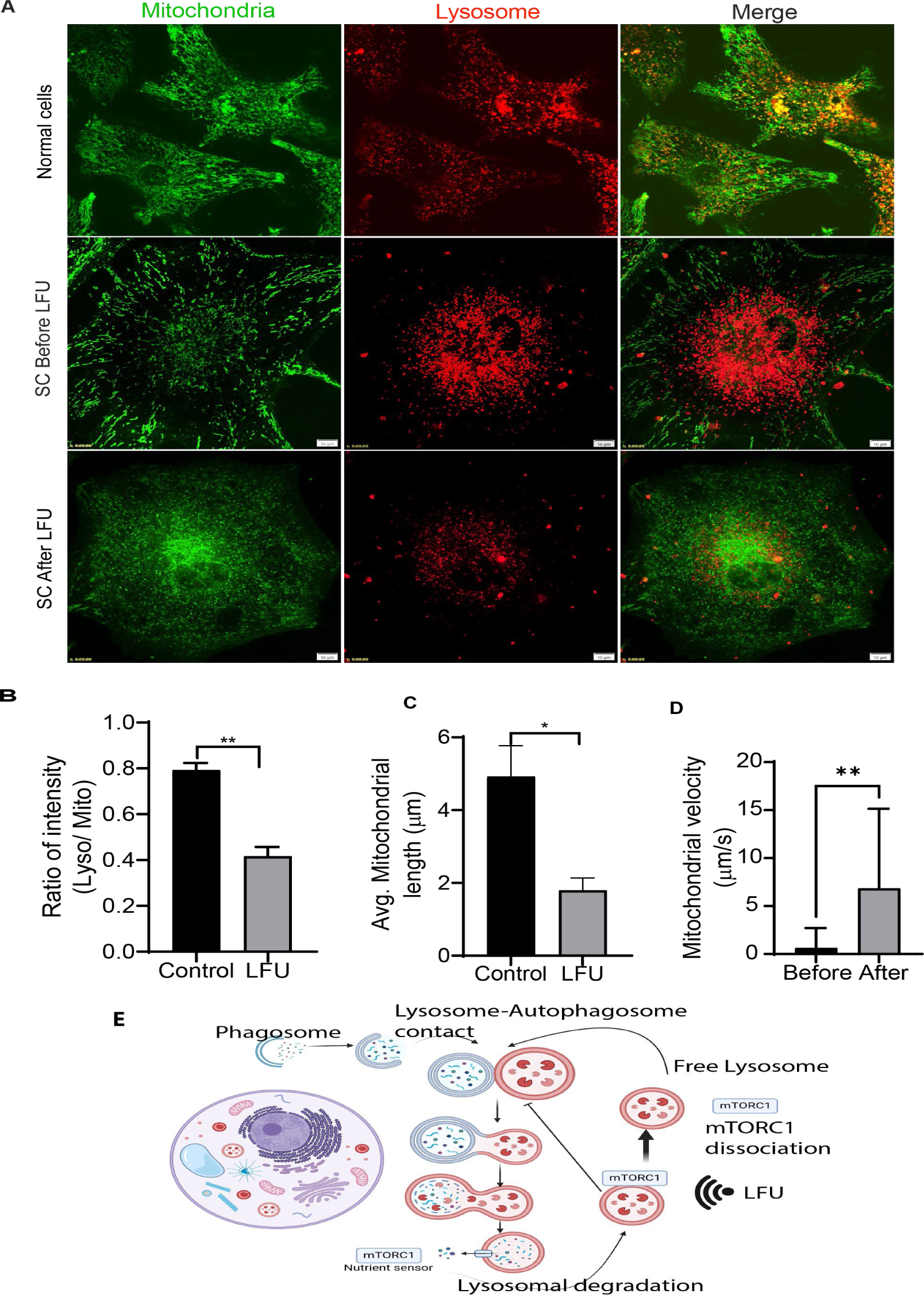
Low Frequency Ultrasound Decreases Mitochondrial Length and Lysosome intensity in Senescent Cells. (A) Representative immunofluorescent images of mitochondrial morphology and lysosome fluorescence in normal, senescent (bleomycin sulfate-treated for 30 hours and incubated for 3 days) and LFU-treated senescent Vero cells stained with Mitotracker and Lysosome tracker. Scale bar=10 µm. (B) Ratio of intensities of lysosomal to mitochondrial staining is decreased by ultrasound treatment of senescent cells. (C) Quantification of mitochondrial length shows decreased length after LFU. Results are shown as mean ± SD, minimally 108 cells were analyzed, n>3 experiments, and significance was determined using two-tailed unpaired t-test. *P<0.05, ** P<0.002. (D) Velocity of mitochondria before and after LFU treatment. Images were capture every five seconds for 10 minutes. A minimum of 10-12 mitochondrial puncta were manually tracking using ImageJ software. (E) Diagram of a working model for the rejuvenation of senescent cells by activation of autophagy through LFU inhibition of mTORC1 activity.

### LFU inhibits SASP secretion

To determine if LFU treatment of SASP-secreting BS-induced senescent cells blocked further secretion of SASP, 24 hour supernatants were collected before and after LFU treatment (Figure S5A). The supernatant from before LFU treatment (S0) inhibited growth and caused an increase in size of normal human foreskin fibroblasts (HFFs) whereas the 24 hour supernatant after LFU (S24) supported normal growth without altering size (Figure S5B, C, and D). In a separate experiment where the levels of SASP components in supernatants from replicatively senescent HFFs were measured before and after LFU treatment, the levels of eight SASP components were lower after LFU exposure, including interleukin (IL)-6, IL-8, IL-10 and IL-15, as well as the pro-inflammatory molecules TNF-α, IFN-γ, VEGF and MIP-1a (Fig. S5E). Thus, it appeared that LFU-mediated rejuvenation of senescent cells blocked the secretion of SASP components.

### LFU of normal cells causes the secretion of components that activate senescent cell growth

Many studies have shown that physical exercise delays peripheral tissue ^50^ and brain ^34^ aging. To test the possibility that LFU could benefit senescent cells through an indirect effect on normal cells, inducing the production of a pro-rejuvenation factor(s), we treated normal HFF cells with LFU over a period of three days (1 h per day) and then collected the supernatant following ultrasound treatment (USS) (Figure 3A). BS-induced senescent HFF cells were treated with USS to determine if a paracrine factor(s) was released from the LFU-treated normal cells that promoted the growth of senescent cells (Figure 3A). Notably, USS promoted the growth of senescent cells (Figure 3B and 3C) and decreased their spread area compared to control normal media-treated senescent cells (Figure 3D). The levels of chemokines and cytokines in the supernatants from the third passage (P3) of non-treated HFFs after 3 days were different from the levels in the USS of P3 LFU-exposed HFFs (USSP) (Figure 3E). The levels of IL-9, IL-17, MCP-1, MIP-1a, MIP-1b, RANTES, GM-CSF, EOTAXIN, G-CSF and IP-10 were lower, while the levels of PDGF-bb and VEGF were higher, in the USSP compared to the control cell supernatants (Figure 3E). Thus, LFU treatment of normal cells stimulated an alteration in the secretion of factors to the supernatant that were associated with growth of senescent cells.

**Figure 3.**
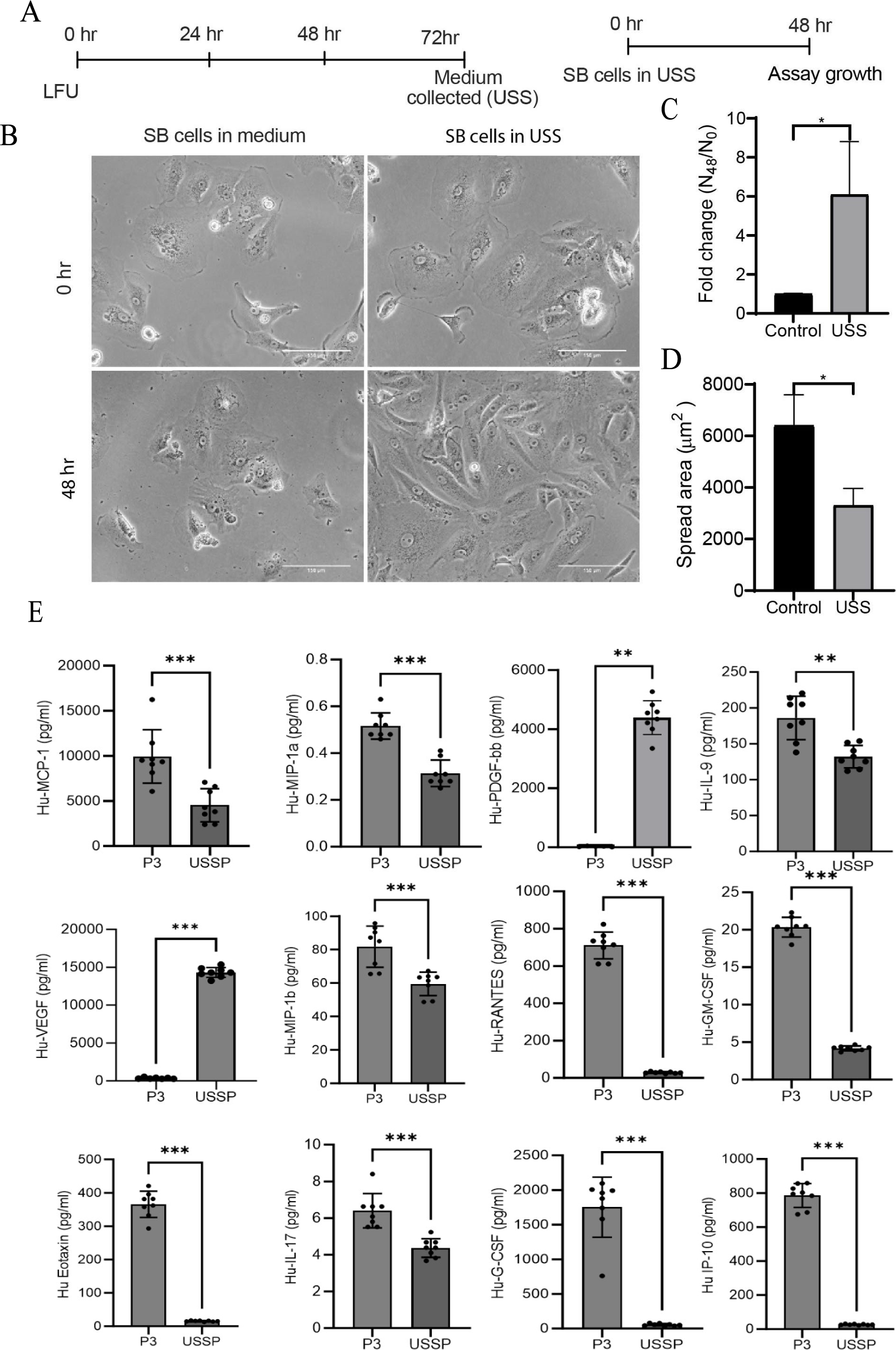
Normal Cells Treated with LFU Secrete Growth Activating Factors. (A) Timeline and strategy of LFU treatment of normal proliferating cells, passage 3 HFF cells. Schematic shows that normal cells were treated with LFU four times in the same media and the supernatant was collected (USS) for incubation with senescent cells for 48 h. (B) Brightfield images show change in morphology of senescent cells in supernatant collected from LFU treated normal cells and after 48 h in the presence of USS. Senescent cells in normal growth medium were the controls. (C) Graph shows growth of SCs in USS increased and (D) shows decreased spread area of senescent cells. (E) Chemokines and cytokines in supernatants collected from untreated and LFU treated cells (P3 HFF) were analyzed using Multiplex immunoassay. Results are plotted as mean± s.d., n=6 replicates. Mann Whitney Test was used to determine the statistical significance, ns not significance, * p value <0.05 ** p value <0.01, *** p value 0.0001, and **** p value 0.00001. Graphs were plotted by mean ± SD; *P<0.05, **P<0.002, *** P<0.0001. Minimum, 200 cells were analyzed from the three independent experiments for graph (C) and (D). Scale bar= 300 µm.

### LFU activates Ca^2+^ entry and autophagy through the mechanosensitive ion channel Piezo1

The first component of a cell that encounters LFU is the plasma membrane. In the case of tumor cell treatment with LFU, Piezo1 ion channels are activated and play a major role in tumor cell apoptosis.^52^ And Piezo1 is known to regulate Ca**^2+^** flux and Ca**^2+^** signaling regulates autophagy.^56,57^ Thus, if LFU is activating growth by activating autophagy downstream of Ca**^2+^** entry, then Ca**^2+^** loading and autophagy activation should occur after LFU treatment. We explored that possibility and found that within 1-2 minutes of the start of LFU, there was a dramatic increase in cytoplasmic Ca**^2+^** that lasted 30-60s (Figure 4A, B). Next, we explored if Piezo1 is the mechanosensitive ion channel regulating LFU-mediated cell growth. We incubated senescent cells with the Piezo1 inhibitor, GsMTx4, and found that it inhibited cell growth after LFU treatment to a level equivalent to GSMTx treatment alone (Figure 4C). Because GSMTx would alter Ca^2+^ homeostasis, it may have significantly increased the growth of the P21 HFFs (Figure 4E). Further, when the senescent cells were treated with LFU in the presence of Ruthenium Red, an inhibitor of many mechanosensitive channels, including Piezo1 ^58^, there was also less growth of senescent cells after 48 h compared to LFU treatment alone (Figure 4D, E). Thus, it appears that LFU stimulation of growth depends upon the activity of the mechanosensitive ion channel, Piezo1.

**Figure 4:**
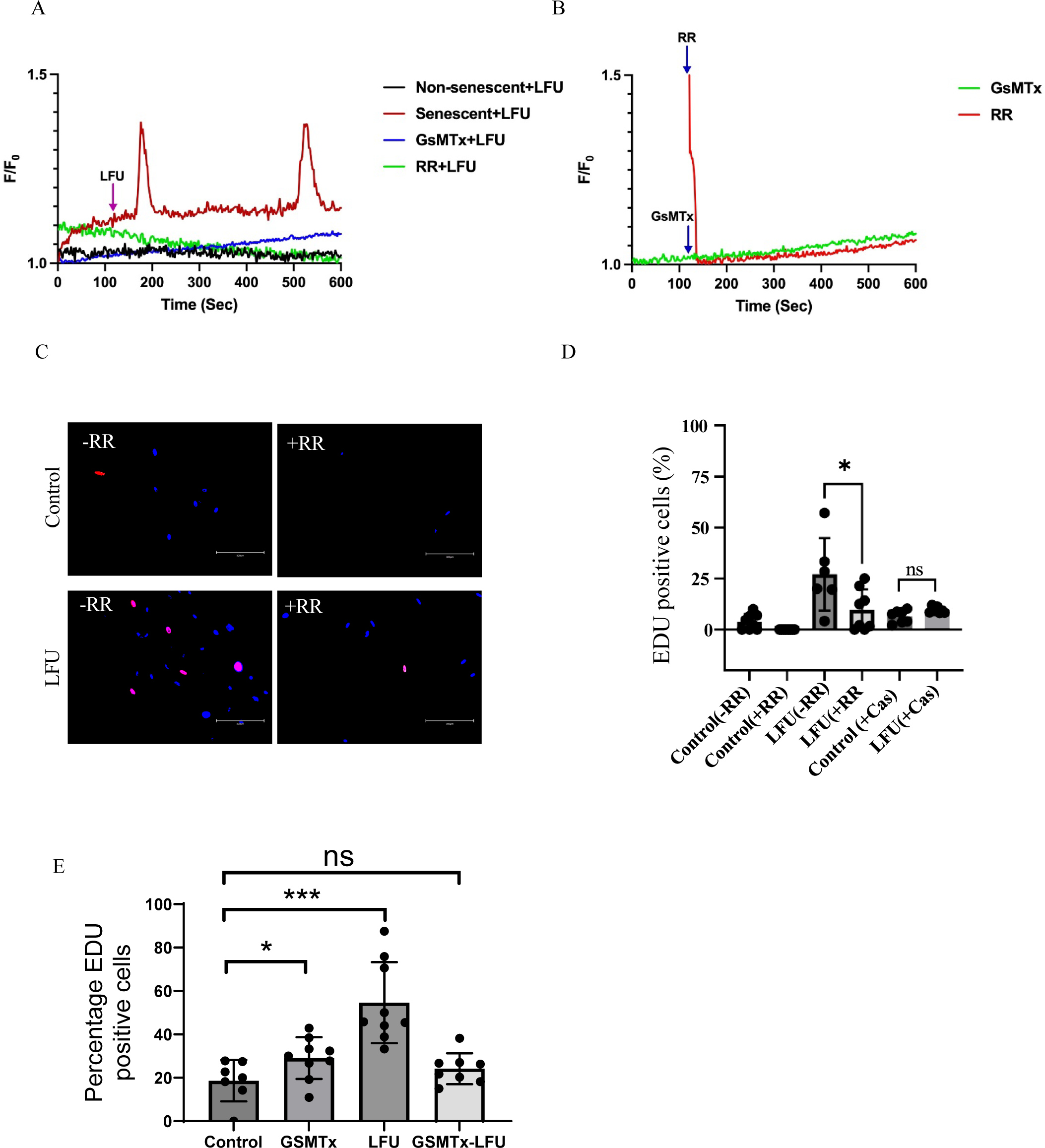
LFU Stimulates Ca2+ entry and Inhibition of Piezo 1 and TRPV1 channels blocks rejuvenation. (A) Time course of Ca^++^ levels in a senescent cells (P21 treated with Piezo1 inhibitor (GDMTx) and TRPV1 inhibitor (Ruthenium Red=RR). (B) Time course of Ca^++^ levels in a senescent cell in presence of Piezo1 and Trpv1 inhibitors. (C) Representative Immunofluorescence-stained images of 3 day old bleomycin-sulfate-induced senescent cells. Cells were treated with or without LFU in presence or absence of TRPV1 inhibitor, Ruthenium Red (RR). Caspaicin (Cas) was used to activate TRPV1(Images are not shown). Scale bar= 300 μm. Cells were then incubated for 12 h in EDU reagents and stained for EDU (red staining) as well as nuclei (DAPI stained, blue). (D), quantification of proliferating cells by manually counting the red colored nuclei. EDU positive cells, and dividing by the total number of cells in the fields.(E) Quantification of cell proliferation of passage 15 HFF cells with or without GSMTx and Ruthenium red. Results are plotted as the mean of three independent experiments ± s.d. * p value < 0.05 using unpaired two tailed test, more than 100 cells were counted in each condition.

Autophagy can be activated by several Ca^2+^-dependent processes ^59^. Thus, we next explored if LFU induces autophagy in senescent cells. We did so by measuring the changes in the GFP-to-RFP fluorescence ratio of a GFP-LC3-RFP construct. When this construct was expressed in senescent cells, the GFP:RFP ratio dropped dramatically after LFU treatment (Figure 5A, B), indicating that GFP-LC3-RFP moved to an acidic compartment that decreased GFP fluorescence indicative of lysosomal processing. We next showed that autophagy is needed for LFU-mediated rejuvenation of senescent cells by showing that inhibition of autophagy by chloroquine diphosphate (CCD) co-treatment blocked LFU-activated autophagy (Figure 5B).

**Figure 5.**
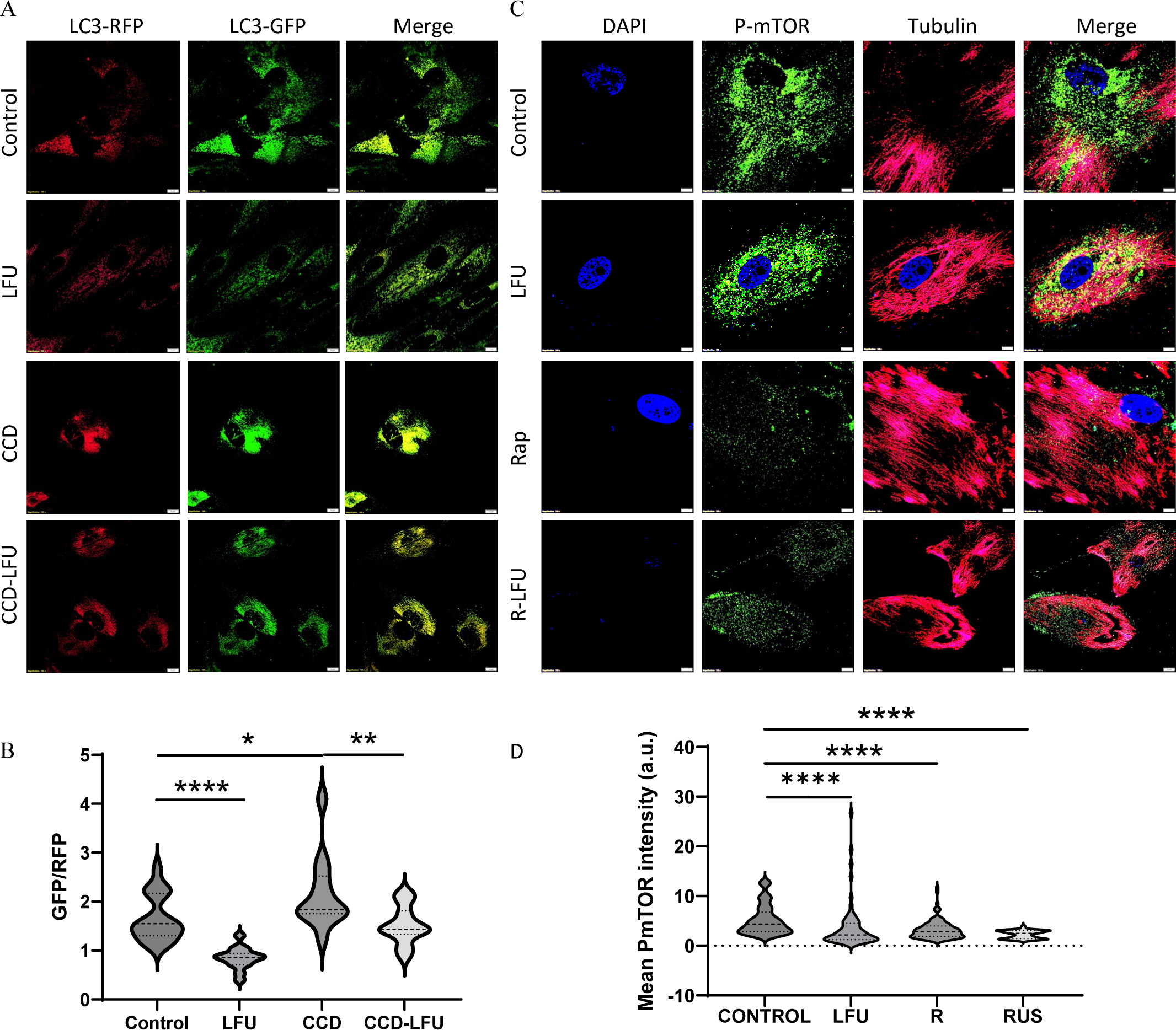
LFU Treatment Increases Autophagy and Decreases mTORC1 Activity. (A) Representative immunofluorescence images of GFP-LC3-RFP transfected in P18 HFF cells treated with LFU with or without Chloroquine diphosphate (CCD, 10 Um). Control cells were not treated with CCD or LFU. Cells were transfected with autophagy sensor, LC3 and incubated overnight as per manufacturer’s instructions (See the method sections for details). Cells were then treated with CCD or LFU and incubated for 24 h. Confocal images were captured after 24 h incubation. Scale bar= 10 um. (B), violin plots show the quantification of relative intensity of GFP to RFP fluorescence. Autophagy flux is inversely correlated with GFP relative intensity because active autophagy quenches the GFP fluorescence. Result is plotted as mean± s.d., n=80 fields ROI and a minimum 10 cells were analyzed. * p-value <0.05, ** p-value <0.01 and **** p value<0.0001, two tailed Mann Whitney test was used. (C), Representative immunofluorescence images of P16 HFF cells (control), (LL) treated L-Leucine (100 mM), (Rap) treated with Rapamycin (1 μM), (LFU) with LFU, (RUS) treated with Rapamycin and LFU, (LUS) treated with LFU and L-Leucine.. Cells were co-immunostained with P-mTORC1 antibody (Green), and Tubulin antibody (Red), and DAPI (Blue). Scale bar= 10 um. (D), violin plots show the quantification of mean intensity of P-mTORC1 (Green) fluorescence. Result is plotted as mean± s.d. per cell, minimum 15 cells were analyzed. **** p value<0.0001 Two tailed Mann Whitney test was used.

### Inhibition of Sirtuin1 blocks LFU-induced rejuvenation

Another protein linked to senescence is Sirtuin1 (SIRT1), an analog of the yeast deacetylase ser1, as it inhibits senescence.^60^ We found that after LFU treatment SIRT1 moved from the nucleus and its level of staining was stronger (Figure S6A and F). We then inhibited SIRT1 with the EX-527 inhibitor during LFU treatment of senescent cells and found that their rejuvenation was blocked and that the SIRT1 levels were lower (Figure S6C, D, E and F). Thus, these data indicate that SIRT1 activity, likely its NAD+-dependent deacetylation function, is needed for rejuvenation effects of LFU on senescent cells.

As LFU caused a relative decrease in GFP fluorescence of the GFP-LC3-RFP construct in the presence or absence of CCD (Figure 5A), we also took a gain-of-function approach. Rapamycin is known to activate autophagy by inhibiting mTORC1 ^43^, and we found that it acted synergistically with LFU to increase growth (Figure S6G) as expected, if LFU inhibition of mTOR was increased by rapamycin (Figure 5C and D). Thus, LFU pressure oscillations are able to reverse the effects of senescence on autophagy as well as mTOR activity.

To further explore the role of SIRT1 in the reversal of senescence via activation of autophagy, we measured the effects of SIRT1 inhibition on the GFP: RFP fluorescence ratio of the GFP-LC3-RFP construct. Addition of a SIRT1 inhibitor, EX-527, either alone or with LFU, caused a significant increase in the GFP:RFP ratio relative to LFU treatment alone, indicating that autophagy was inhibited (Figure S6B). Thus, SIRT1 activity is needed for the strongest activation of autophagy during LFU treatment.

### Various readouts of cell stress are restored to normal by LFU treatment of senescent cells

To next explore the effect of LFU on senescence, we tested for several readouts of cell stress that occur during senescence. Notably, we found that in BS-induced senescent Vero cells there were significantly lower levels of p53 staining and γH2AX foci, as well as the levels of H3K9me3, ROS and MitoROS, after LFU treatment compared to control-treated senescent cells (Figure S7A-H). Thus, by many criteria, LFU treatment caused a dramatic decrease in the senescent state.

Other factors have been linked to aging and cell senescence, included telomere shortening ^61^ and DNA hypomethylation.^62^ In the case of telomere length, there was a striking increase in the average telomere length of replicatively senescent HFF fibroblasts after LFU treatment (Figure S7I) and a small increase in replicatively senescent mesenchymal stem cells (MSCs) (Figure S7J). We also found that the DNA methylation levels of replicatively senescent HFF cells were higher after LFU treatment (Figure S7K, L), which is consistent with the rejuvenation effect induced by LFU in these cells and with the decrease in H3K9me3 levels after LFU (Figure S7D, E). Thus, LFU treatment of senescent cells reverses senescence-dependent changes in the nucleus, consistent with a restoration of the normal cell state by LFU.

### RNA-seq analysis reveals major transcriptional changes after LFU

We next performed RNA-seq analysis of LFU-treated senescent cells and found that the expression levels of 50 genes were upregulated by a factor of two or more upon rejuvenation whereas 140 genes were downregulated (Tables 1, 2). Of particular interest were the SASP proteins that were downregulated upon rejuvenation, which included IGF2, IGFBP2, FGF7 and C1Q-TNF7. We performed further pathway analysis and found that many of the genes activated by LFU were in pathways induced by viral infections (Tables 3, 4). This is consistent with the activation of cell growth upon rejuvenation.

**Table 1.** RNA-seq Data.

RNA-seq Data
| **List of Upregulated Genes in LFU-Rejuvenated HFFs vs Senescent Cells** |  |  |  |
| --- | --- | --- | --- |
| **S. No.** | **Name** | **log<sub>2</sub>FoldChange** | **P-value** |
| 1 | IFI27 | 4.706732879 | 3.39E-139 |
| 2 | ANO3 | 4.303317562 | 7.94E-19 |
| 3 | BST2 | 4.267516399 | 4.60E-48 |
| 4 | NGFR | 4.227721006 | 3.99E-15 |
| 5 | OAS1 | 4.194394186 | 2.58E-109 |
| 6 | OAS2 | 3.851520822 | 3.19E-205 |
| 7 | IFI6 | 3.687238684 | 0 |
| 8 | NTSR1 | 3.587703179 | 5.84E-20 |
| 9 | MX1 | 3.41456665 | 7.56E-188 |
| 10 | MX2 | 3.353896447 | 4.53E-85 |
| 11 | IFITM1 | 3.267590391 | 1.12E-82 |
| 12 | SFRP1 | 3.062130153 | 1.02E-136 |
| 13 | OAS3 | 3.039643219 | 3.46E-89 |
| 14 | RSAD2 | 3.00663542 | 9.46E-26 |
| 15 | CMPK2 | 2.992910736 | 4.73E-32 |
| 16 | EDARADD | 2.971533613 | 5.08E-10 |
| 17 | GBX2 | 2.893735519 | 9.38E-11 |
| 18 | ENSG00000225886 | 2.869460744 | 7.25E-07 |
| 19 | NEURL3 | 2.844376063 | 2.79E-05 |
| 20 | SLC38A11 | 2.839565113 | 1.67E-59 |
| 21 | SPHKAP | 2.79074685 | 1.05E-25 |
| 22 | ESM1 | 2.774304098 | 1.52E-48 |
| 23 | HAS2 | 2.761341752 | 1.28E-48 |
| 24 | HSPA6 | 2.608245534 | 3.99E-10 |
| 25 | CCL5 | 2.545666491 | 1.38E-20 |
| 26 | HERC6 | 2.543077514 | 5.23E-87 |
| 27 | LINC01234 | 2.475565502 | 4.77E-08 |
| 28 | ANGPT2 | 2.472972104 | 2.32E-05 |
| 29 | TMEM158 | 2.442808274 | 2.10E-59 |
| 30 | DIRAS3 | 2.426571346 | 4.54E-09 |
| 31 | IFI44 | 2.382909102 | 1.83E-78 |
| 32 | EPSTI1 | 2.376583283 | 1.42E-66 |
| 33 | ZNF385D | 2.372390644 | 4.42E-10 |
| 34 | STAC | 2.368386566 | 1.66E-11 |
| 35 | GBX2-AS1 | 2.334084241 | 5.12E-07 |
| 36 | GNA15 | 2.318072578 | 2.24E-06 |
| 37 | IFI44L | 2.303837053 | 5.65E-37 |
| 38 | HERC5 | 2.285450475 | 4.55E-44 |
| 39 | COLEC12 | 2.264077281 | 2.70E-15 |
| 40 | ISG15 | 2.259660967 | 5.02E-145 |
| 41 | ANO9 | 2.258562492 | 7.43E-05 |
| 42 | FRMD3 | 2.243609765 | 1.57E-27 |
| 43 | CD36 | 2.218190589 | 1.51E-15 |
| 44 | IFIH1 | 2.192859031 | 1.26E-42 |
| 45 | HAND2 | 2.127703818 | 3.13E-19 |
| 46 | BLK | 2.055308732 | 3.31E-05 |
| 47 | IFIT1 | 2.051756712 | 1.68E-146 |
| 48 | USP18 | 2.047027284 | 8.39E-32 |
| 49 | SLC24A3 | 2.035592849 | 1.77E-45 |
| 50 | OASL | 2.029842409 | 1.58E-21 |
| 51 | LRRC37A6P | 2.02025571 | 1.49E-06 |
| 52 | PODXL | 2.016215331 | 1.31E-35 |
| 53 | EPB41L3 | 2.000835065 | 5.10E-127 |

**Table 2:**
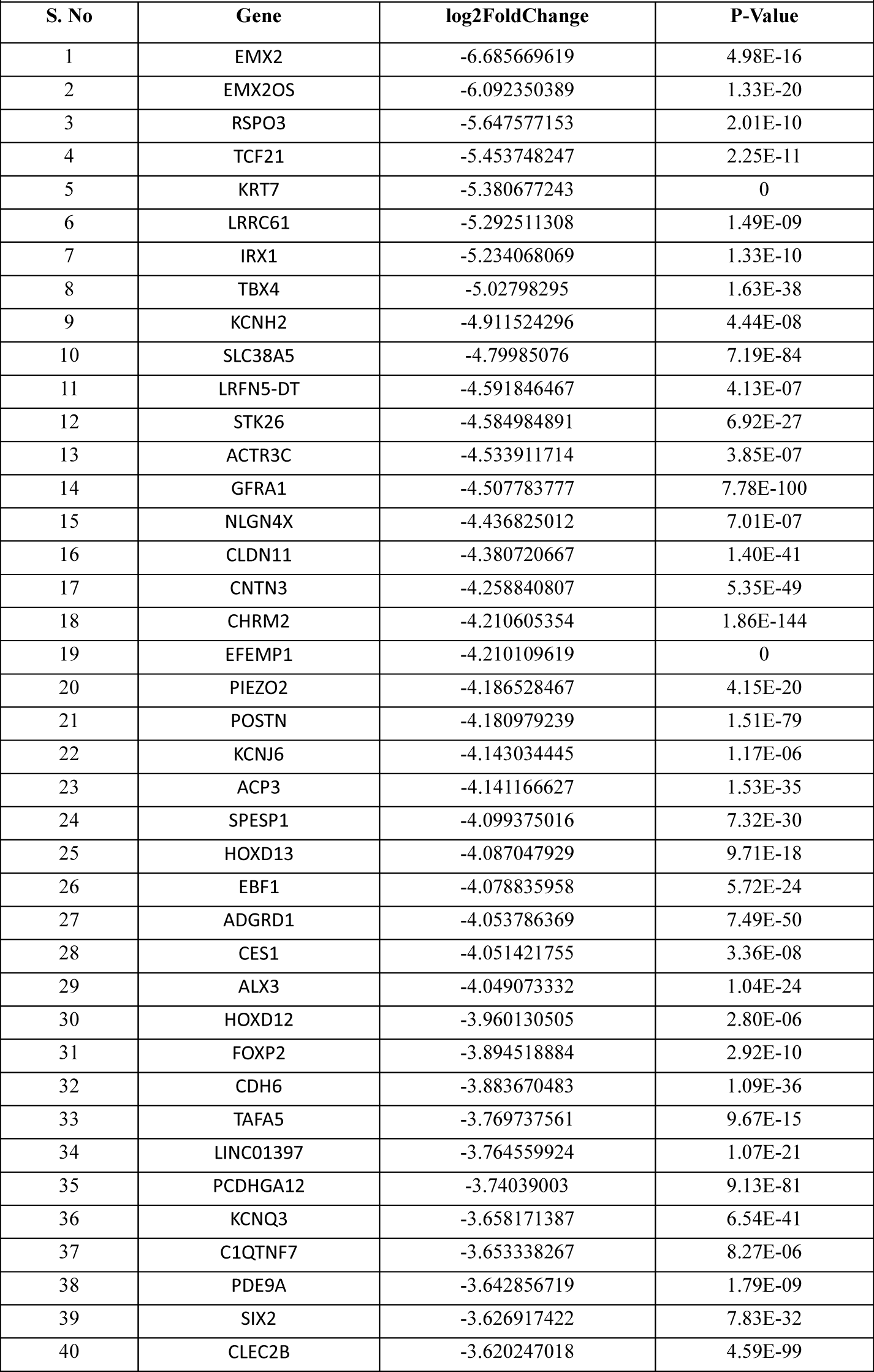
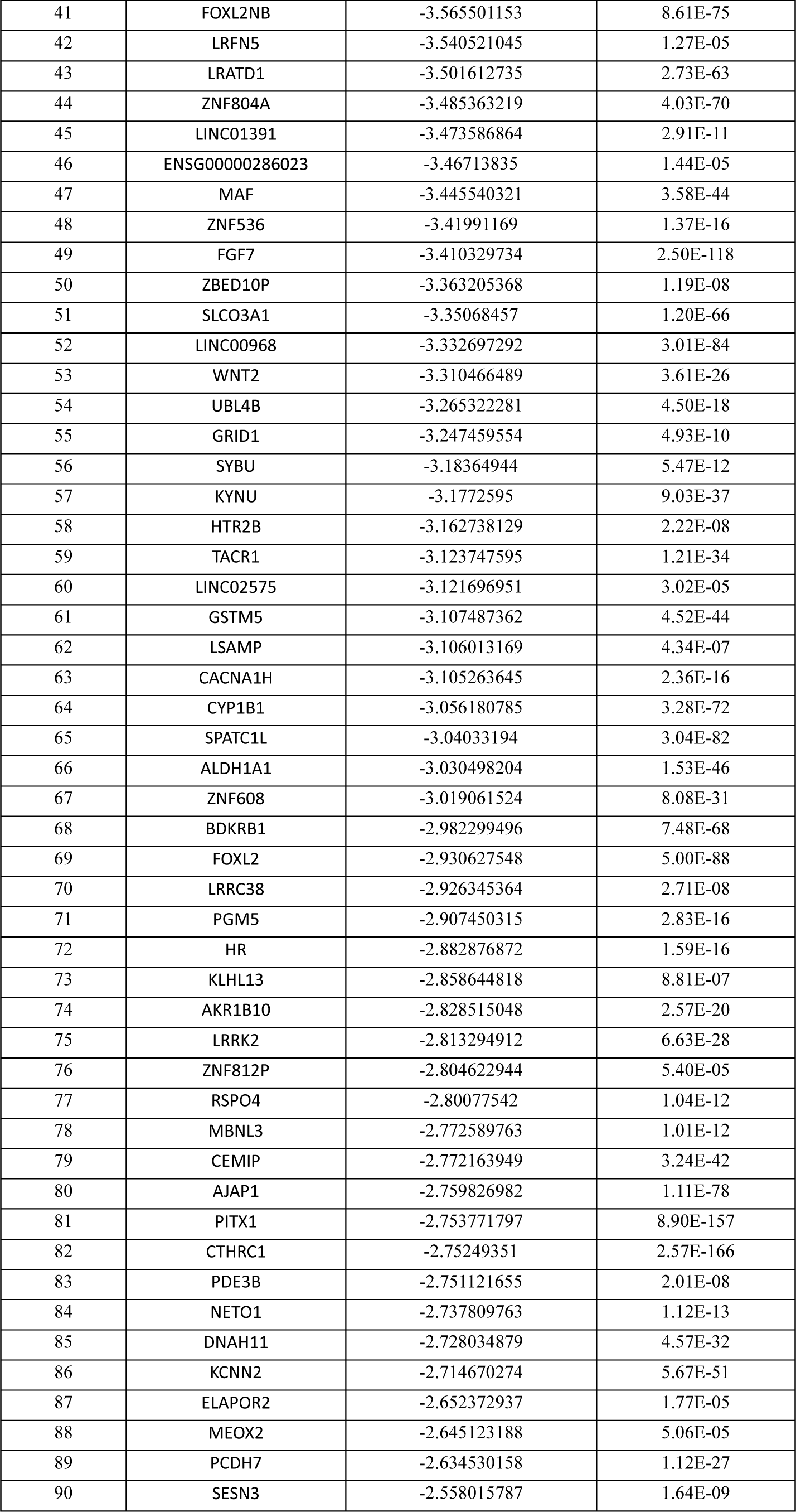
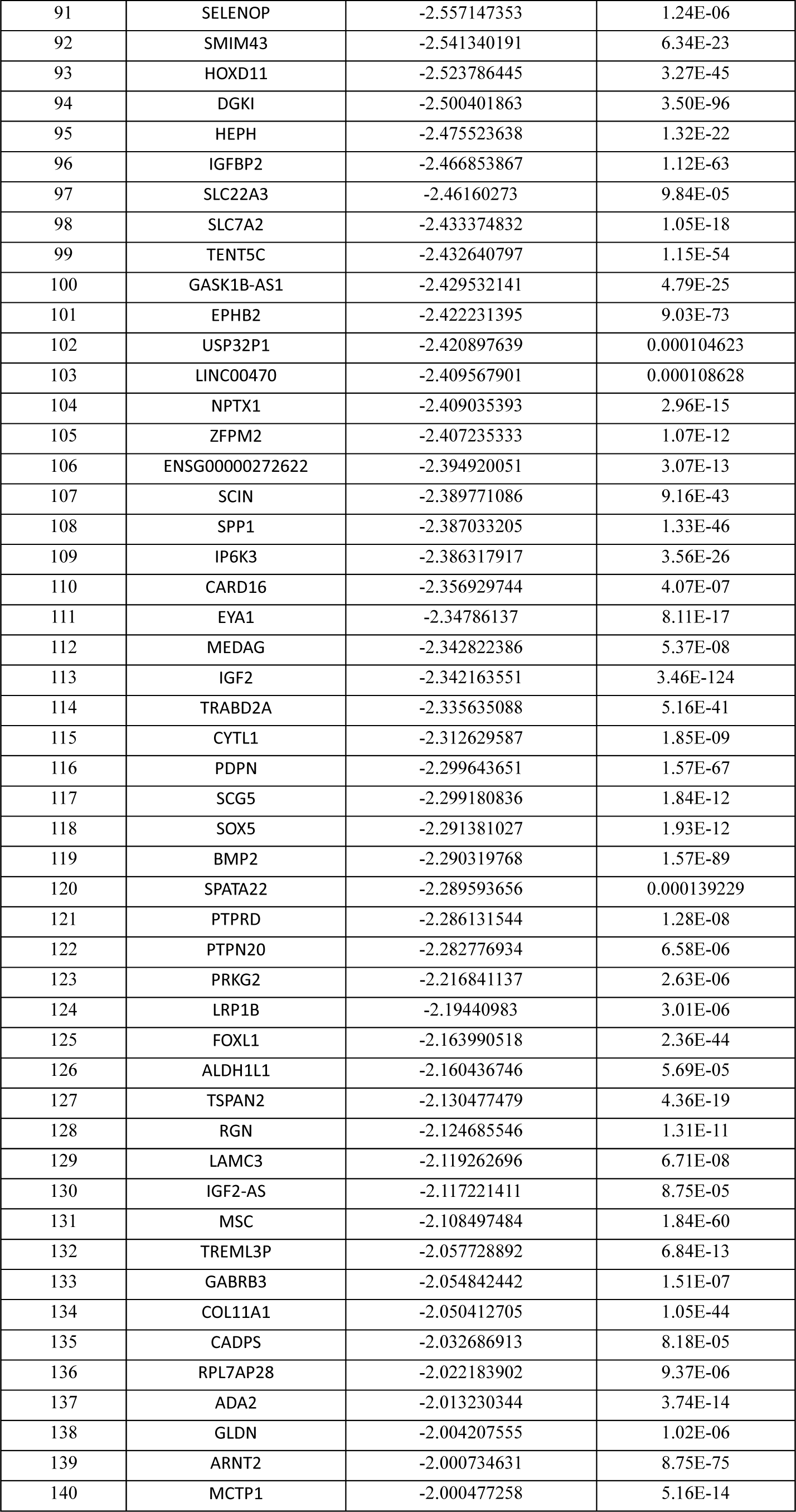
List of Downregulated Genes in LFU-Rejuvenated HFFs vs Senescent Cells.

**Table 3:** List of Genes upregulated in LFU-treated late-passage HFFs.

| Bio Process | p-value | q-value | overlap_genes |
| --- | --- | --- | --- |
| Influenza A | 1.619929e-08 | 9.881566e-07 | [IFIH1, RSAD2, OAS1, OAS2, MX2, OAS3, CCL5, MX1] |
| Measles | 7.942201e-08 | 2.422371e-06 | [IFIH1, OAS1, OAS2, MX2, OAS3, MX1, HSPA6] |
| Hepatitis C | 1.828574e-07 | 3.718100e-06 | [RSAD2, OAS1, OAS2, MX2, OAS3, MX1, IFIT1] |
| Coronavirus disease | 2.527530e-06 | 3.854483e-05 | [IFIH1, OAS1, OAS2, MX2, OAS3, MX1, ISG15] |
| NOD-like receptor signaling pathway | 1.344502e-03 | 1.640292e-02 | [OAS1, OAS2, OAS3, CCL5] |
| Herpes simplex virus 1 infection | 1.970480e-03 | 1.750871e-02 | [BST2, IFIH1, OAS1, OAS2, OAS3, CCL5] |
| Epstein-Barr virus infection | 2.009196e-03 | 1.750871e-02 | [OAS1, OAS2, OAS3, ISG15] |
| Human papillomavirus infection | 1.141362e-02 | 8.702885e-02 | [MX2, MX1, ISG15, OASL] |
| RIG-I-like receptor signaling pathway | 1.483205e-02 | 1.005283e-01 | [IFIH1, ISG15] |
| Lipid and atherosclerosis | 1.935232e-02 | 1.180492e-01 | [CCL5, HSPA6, CD36] |

**Table 4:** List of Genes downregulated in LFU-treated late-passage HFFs.

| Bio Process | p-value | q-value | overlap_genes |
| --- | --- | --- | --- |
| GnRH secretion | 0.001055 | 0.119259 | [KCNJ6, SPP1, KCNN2, CACNA1H] |
| Calcium signaling pathway | 0.006934 | 0.309739 | [CHRM2, FGF7, HTR2B, BDKRB1, TACR1, CACNA1H] |
| Serotonergic synapse | 0.008223 | 0.309739 | [GABRB3, KCNJ6, HTR2B, KCNN2] |
| Regulation of actin cytoskeleton | 0.018747 | 0.394402 | [CHRM2, ACTR3C, FGF7, SCIN, BDKRB1] |
| GABAergic synapse | 0.024628 | 0.394402 | [GABRB3, KCNJ6, SLC38A5] |
| Morphine addiction | 0.026083 | 0.394402 | [GABRB3, KCNJ6, PDE3B] |
| Circadian entrainment | 0.030721 | 0.394402 | [KCNJ6, PRKG2, CACNA1H] |
| Neuroactive ligand-receptor interaction | 0.033034 | 0.394402 | [CHRM2, GABRB3, GRID1, HTR2B, BDKRB1, TACR1] |
| Pathways in cancer | 0.033440 | 0.394402 | [ARNT2, FGF7, BMP2, LAMC3, IGF2, BDKRB1, WNT2, GSTM5] |
| Tryptophan metabolism | 0.034903 | 0.394402 | [KYNU, CYP1B1] |

### Rejuvenation is increased by Rho kinase inhibition

Previous studies from our laboratory reported that a different ultrasound treatment caused apoptosis of tumor cells but not normal cells.^52,53^ In that case, LFU-dependent tumor cell apoptosis increased after microtubule depolymerization by nocodazole and the Rho kinase inhibitor (Y-27632) blocked apoptosis. Depolymerization of microtubules increases Rho kinase activity and myosin contractility ^55^ whereas the Rho kinase inhibitor decreases contractility. Thus, we co-treated senescent HFF cells with LFU and Y-27632 and found greater rejuvenation than the LFU alone control while microtubule depolymerization with nocodazole had no effect (Figure S6H). Thus, the reversal of senescence by LFU is different from the activation of tumor cell apoptosis, as they involve different cytoskeletal elements.^55^

### Expansion of replicative senescent cells by LFU

Because replicative senescence is presumably a common property of normal cells that limits their growth ^3^, we then tested whether LFU could extend the growth potential of normal cells. After 15 passages HFFs showed a slower growth rate, a greater average cell size and exhibited increased β-galactosidase activity (Figure 6A, B and C). However, after LFU treatment with every other passage, for passages 15-24, HFFs behaved like normal cells and continued to grow without a significant change in growth rate for at least 24 passages (Figure 6A). This made it possible to grow >7,500 (2^13^)-fold more HFFs than without LFU, while the spread area was normal (Figure 6B) and there was no increase in the percentage of cells expressing β-galactosidase (Figure 6C). When P24 LFU-treated cells were cultured on soft matrices, they ceased to grow, showing rigidity-dependent growth. Similarly, we expanded MSCs with LFU beyond their normal replicative limit (Figure 6D). Upon treatment with differentiation medium, the LFU-expanded P18-MSCs differentiated into adipocytes or osteocytes, depending upon the medium (Figure 6E-G) although the level of adipocyte differentiation was higher in the LFU-treated cells than in the replicatively senescent MSCs.

**Figure 6.**
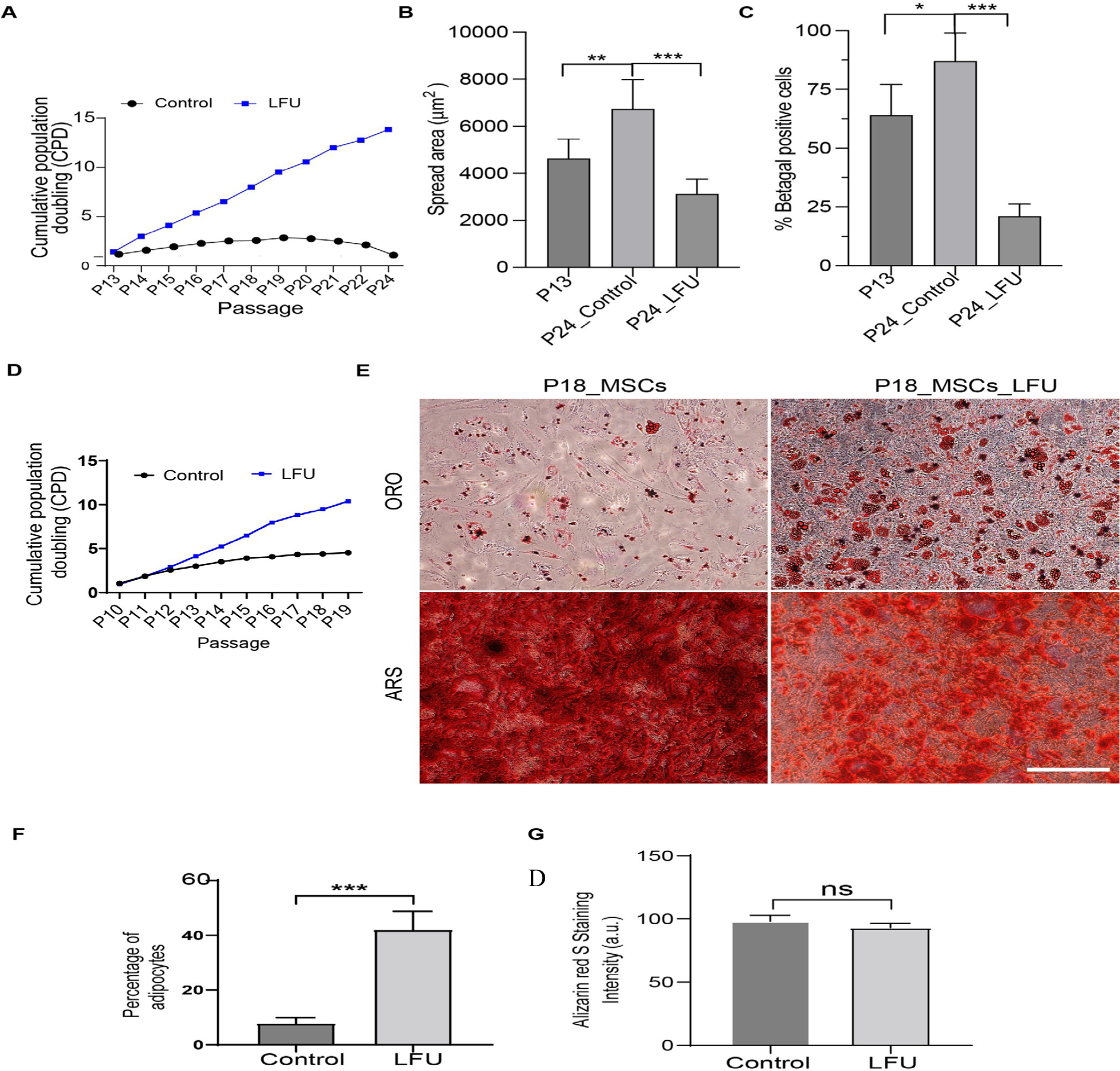
Ultrasound Reversal of Replicative Senescence Increases Number of Cells. (A) Growth rate shown as cumulative population doublings (CPD) for Control HFF and LFU treated HFF cells passaged every 48 h from P13 to P24 passage with LFU treatment every other passage (B). LFU treated cells were smaller than the p24 control and even p13 cells. (C) Percentage of SA-b-galactosidase positive cells decreased after LFU treatment. (D) Similarly, LFU treatment of mesenchymal stem cells (MSCs) expanded the cell number P10-P19, with treatment every other passage. (E) LFU treated MSCs showed normal differentiation to (ORO) adipocytes and (ARS) osteocytes. Oil red o staining dye marked lipid droplets (ORO) and alizarin red S-stained dye marked osteogenesis (ARS). (F) Percentage adipocytes were quantified in P18 MSCs treated with and without LFU. Results are plotted as mean of three independent experiments, a minimum of 100 cells were counted. (G) Quantification of osteocytes was determined by the intensity of alizarine Red S staining. Mean intensity was calculated from ten random images of three independent experiments. Results are shown as mean ± SD, minimally 200 cells for spread area and 150 cells for percentage b-galactosidase analysis, n>3 experiments, and significance was determined using two tailed unpaired t-test. . *P<0.05, **P<0.002, *** P<0.0001.

### The mouse healthspan and lifespan is improved by LFU in a dose-dependent manner

To determine if LFU can be utilized *in vivo* to improve the performance of older mice, we followed one sham-treated (placed in the water bath without LFU for 30 min every day) and 5 LFU-treated groups of mice (*n* = 10 mice per group), with the 5 LFU-treated groups given a different dose schedule (either daily (D1), every other day (D2), or every third day (D3) with 1X power and daily with 1.3X (1.3X) or 2X (2X) power levels for 30 min) (Figure S8A). After two treatment periods of two weeks each, with a two-week break between treatment periods, we found that physical performance, as measured by either the duration of clinging or of running, improved with all LFU conditions and was statistically significant for all groups in the treadmill and 3 of 5 in the inverted cling test (Figure S8B-E). While LFU treatment for 4 weeks appeared to be more effective than treatment for 2 weeks there was no clear difference among the different regimens within each time frame of treatment.

We sacrificed two mice from each group 5 days after the last LFU treatment and inspected their kidneys and pancreas (Figure S9A). After sectioning, fixing and staining for β-galactosidase, there was ∼70% of the kidney area and ∼70% of the pancreas area showing staining in the sham animals (Figure S9B-E). In contrast, sections of these organs from the LFU-treated animals had only 10-20% of the area stained for β-galactosidase, depending upon the treatment regimen (Figure SB-E). Overall, LFU-treated animals had a significantly smaller fraction of cells in these two tissues expressing the senescence marker, β-galactosidase, which was indicative of their improved performance.

To test the effect of LFU on lifespan of aged mice, we continued to treat the 46 remaining mice <u>in</u> the six groups outlined above for over 300 days (some mice reached 3 years of age) and we added 4 mice that were kept in a cage the whole time (negative control group). In the five LFU-treated groups, the best survivors were those that were given the lowest doses of ultrasound (D2 and D3) with a combined value of about 50% survival at 1000 days (∼33 mo of age) (Figure 7A, B). The best survivors were in the D2 cohort of 7 mice treated every other day with 1X LFU and 3 of those mice survived until 3 years (> 40%). Thus, there was statistically significant evidence that LFU treatment increased longevity but the small number of animals per group (7 or 8) did not allow for statistical significance for individual treatment regimens.

**Figure 7.**
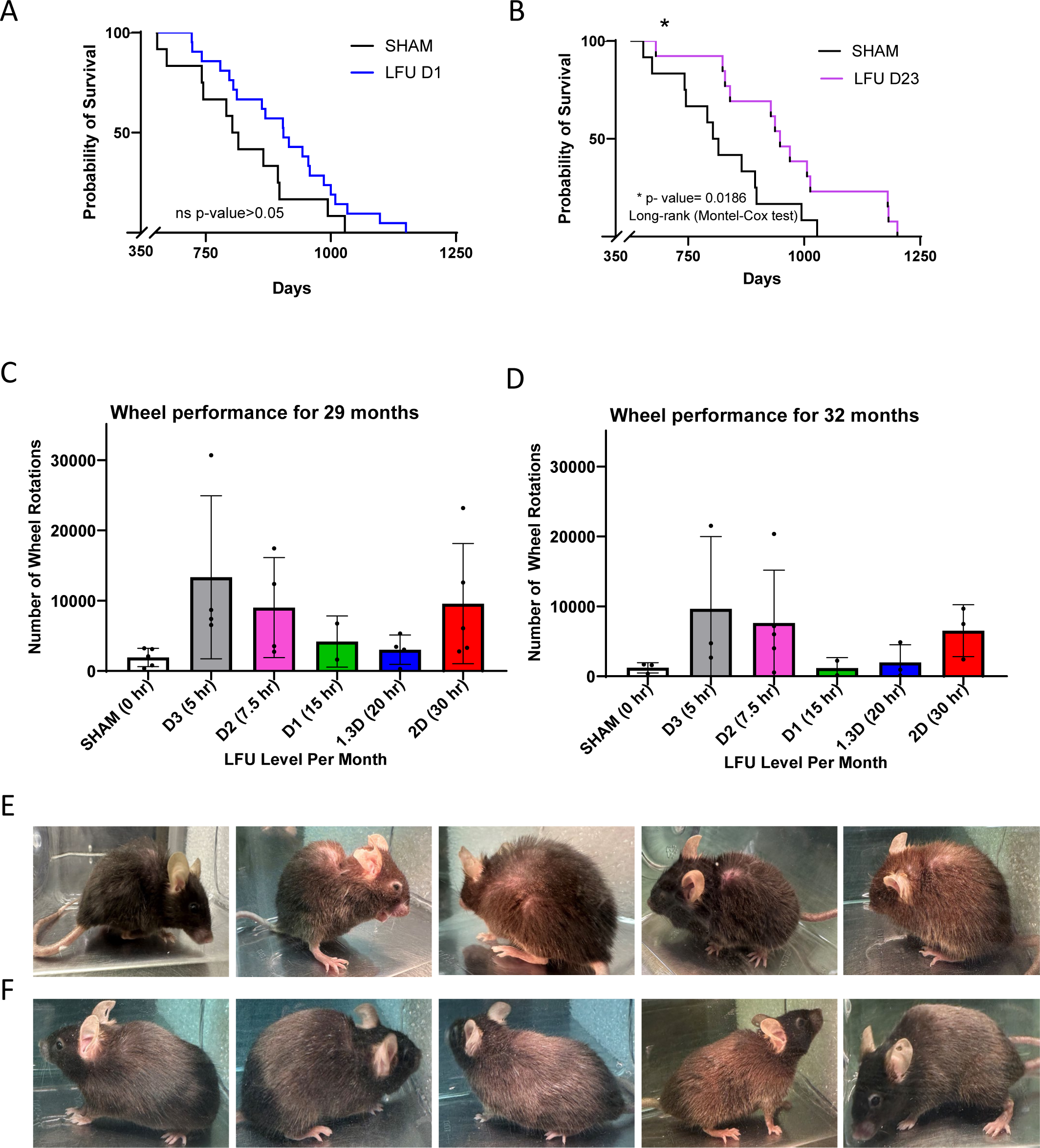
LFU significantly improves the physical performance and lifespan of old mice. (A), survival curves showing the sham and negative control (11 mice) and the LFU treated mice (8-D1, 7-1.3X and 6-2X for a total of 21 mice). *p value <0.05. (B), Survival graphs showing percentage survival curves of the sham and negative control (11 mice) and LFU treated mice (6-D3 and 8-D2). (C and D), Bar graphs showing mouse wheel 5 running activity of sham, and all LFU treated cohorts for mice C) 29 months and D) 32 months of age. Mice were treated with various LFU doses and then placed in the wheel cages. Wheel activity was measured for three days. Results are plotted as mean± s.d. Sham n=6 and LFU treated mice n= 6-8, * p value <0.05 using two tailed unpaired t-test. (E), Pictures showing the side view of sham and (F), 2X LFU treated mice at 30 months of age. There is a video of a representative sham and a 2X mouse at 30 months that further illustrates the difference in activity of the mice (Videos 5 & 6, respectively).

The differences in the activity levels of the sham and treated groups were very striking as seen by videos of their movements (video 5) and by their spontaneous turning of a wheel in their cage (Figure S7C, D). The videos show that two representative sham-treated mice at 29 months of age were slow in their movements and had poor fur density with a balding spot on their backs (videos 5 and 6), whereas two representative mice of the 2X power-treated group were much more attentive and had a denser and darker fur coat (videos 7 and 8). These results are indicative of other members of their respective groups (Figure 7E, F). When the spontaneous activity of each group was measured in a cage with a wheel for 3 days, the LFU-treated animals were particularly active in the D2 and D3 groups and the level of activity correlated with their survival. At 29 months of age, the sham mice turned the wheel only 1500 cycles on average with the best performers at 2000 cycles, whereas the D3 LFU-treated mice had an average of ∼12,000 cycles with the best at 30,000 (Figure 7C). D2 mice turned the wheel at 9,000 cycles on average (Figure 7C) and these two groups had the highest survival levels (Figure 7B). Three months later, the performance of D3 dropped to 10,000 cycles and D2 to ∼8,000 cycles, whereas the sham mice were at ∼1000 cycles (Figure 7D). Thus, the activity levels of the LFU-treated mice were 7-10 fold greater than the activity levels of the sham mice; whereas the survival levels were greater on average and correlated with activity.

In terms of the safety of LFU, half of the mice were treated daily with LFU for over 300 days without damage or evidence of harm from the LFU treatment. Further, the treated mice maintained a normal weight and the animals that died had no tumors or obvious cause of death upon autopsy (data not shown). Thus, there was no evidence of damage by LFU and the highest dosage of LFU (2X) had the effect of keeping the fur thick and dark even at 29 months of age (Figure 7F); whereas the sham controls had a bald spot on their backs and a lower fur density (Figure 7E). This strongly supports the hypothesis that LFU rejuvenated the skin and hair cells, enabling them to produce fur like much younger animals.

## Discussion

Here, we show that senescent cells, as rigorously defined by many markers, including the expression of β-galactosidase, can be mechanically rejuvenated by LFU without transfection or other biochemical manipulations. The ultrasound pressure waves restore normal behavior irrespective of whether senescence is induced by chemical treatment or by repeated replication. There is no apoptosis with LFU and videos of senescent cells before and after LFU show a dramatic increase in cell and mitochondrial motility, as well as in growth. Many features of senescent cells are all reversed by LFU, including the increase in β-galactosidase activity, p21 expression, decreased telomere length, increased H3K9me3 levels, decreased 5mc levels, increased cell size, secretion of SASP and inhibition of growth. Restoration of normal behavior correlates with a decrease in mitochondrial length and lysosomal volume. We also optimized the values of LFU power, frequency and duty cycle to best induce such rejuvenation of senescent cells that belie an unknown set of processes that are mechanically activated by LFU. Surprisingly, ultrasound treatment of normal cells causes secretion of growth stimulating factors that partially restore normal behavior in senescent cells. Because replicative senescent cells are restored to a normal phenotype by LFU, they can be cultured for long periods to produce increased numbers of cells without major alteration in their phenotype.

It is perhaps surprising that fully senescent cells can be rejuvenated by pressure waves. This raises the question of how a senescent cell is defined. Cells that were made senescent by toxic compounds or repeated replications were incubated for long periods and time lapse video microscopy verified the absence of any growth. After such treatments, quiescent cells were not present as over 95% of the cells expressed β-galactosidase ^54^ and many of the larger senescent cells grew and divided in the videos after LFU. By tracing individual cells, we were able to determine that growth was occurring in over 30% of the originally non-dividing cells after 4-5 days. Such robust growth is inconsistent with the growth occurring in a subpopulation of cells that were not senescent. Further, there was no apoptosis after LFU treatment of the senescent cells and over fifteen characteristics of senescence were reversed. Thus, given that all these objective criteria indicate that LFU reversed senescence, we suggest that LFU actually rejuvenates senescent cells. This opens many new possibilities in the aging research field, including the possibility of rejuvenating aged cells *in vivo* to inhibit age-dependent disorders, which appears to be the case based on the results of our mouse studies in this report.

The selective lysis of senescent cells is an alternative approach to reducing the effects of aging, and it has been shown to improve performance of older mice.^8,35,63,64^ The obvious difficulty is that the loss of senescent cells is hard to reverse and the stimulation of growth of the remaining non-senescent cells will stress them and encourage their senescence. In our longevity studies, LFU treatment increased lifespan and the physical performance of aged mice. Thus, it seems that LFU can be used to rejuvenate aged animals without the use of senolytics.

Mechanical effects on cell behavior have been known for a long time. However, recently it has become clear that controlled mechanical perturbations can reproducibly alter cell functions and phenotypic behaviors. Tumor cells are mechanosensitive as either stretching, fluid shear or ultrasound can cause apoptosis *in vitro.*^52,65,66^ In addition, exercise appears to inhibit tumor growth *in vivo.*^67^ Normal cells appear to do better with exercise and myokines that are released with exercise benefit the organism. In the studies presented here, LFU stimulation of normal cells causes the release of beneficial factors that stimulate growth of senescent cells and perhaps could augment the LFU-mediated rejuvenation of aged cells in tissues.

There is evidence of a correlation between exercise and a reversal of senescent cells in older animals and humans.^41^ Individual cells have not been followed in such studies and it is not clear if exercise reverses senescence or is a senolytic.^17^ Here, it is clear that ultrasound pressure waves alone can reverse senescent cell behavior to that of normal cells; *i*.*e*., rejuvenate them, without causing cell death *in vitro*. We suggest that both exercise and LFU will rejuvenate cells *in situ* without apoptosis and thereby increase the performance of aged animals. Because LFU can easily penetrate the whole human body with only significant loss of power in bone and lung, it can rejuvenate most of the tissues, including internal organs, such as the pancreas and kidney, that are not be particularly sensitive to exercise. In this way LFU may have advantages over exercise to improve healthspan and possibly lifespan. But whatever the intervention is, the critical issue for improving the performance of aged animals is the need to decrease the level of SASP and other inhibitory factors, whether it is through the use of senolytics, exercise or LFU, or a combination.

The senescent cell state has been extensively studied but the molecular bases for the changes are not fully understood. It is noteworthy that Ca^2+^ transients occur soon after LFU stimulation and Ca^2+^ is needed for activation of autophagy (Bootman et al., 2018). In addition to autophagy, there are major roles for changes in mitophagy in senescence.^39,43,60,68–70^ Accordingly, most models of the senescence process postulate complicated roles for autophagy and mitophagy to couple changes in activity of lysosomes, mitochondria and other cellular organelles with the cell cycle.^71,72^ The subcellular effects of ultrasound appear to be mediated by Ca^2+^ effects on mechanically-dependent mitochondria-ER-lysosome interactions, which activate lysosomal autophagy and mitochondrial fission that is a requisite for mitophagy. Thus, we suggest that LFU-induced physical distortions act on organized elements of the cytoplasm, like exercise, to reverse molecular complexes induced by aging. Similarly, the increase in motile activity caused by LFU indicates that there is a general effect on cells to restore normal functions that are slowed in senescent cells by the accumulation of damaged material. At a molecular level, senescence is associated with active mTORC1 binding to lysosomes thereby inactivating autophagy.^73,74^ Rapamycin inhibition of mTORC1 is synergistic with LFU-induced rejuvenation and rejuvenation involves activation of autophagy and a decrease in lysosomal staining. In the case of SIRT1, loss of activity is associated with an increase in senescence, which fits with the need for SIRT1 activity in rejuvenation through increased autophagy.^60^ In addition, AMPK plays a major role in mitophagy and autophagy and its activation by AICAR increases the performance of aged mice.^75^ We show here that Piezo1 ion channel activity is needed for optimal LFU rejuvenation of senescent cells and appears to have a role in endothelial cell aging.^76^ The Ruthenium red inhibitor may have effects on multiple targets and selective inhibitors are needed to understand how the relatively slow Ca^2+^ waves might be involved in rejuvenation. In general, much more research is needed to understand the detailed mechanisms by which LFU-induced pressure waves activates autophagy, inhibits mTORC1 and activates SIRT1 function to stimulate cell growth.

Cellular changes with senescence are extensive and involve not only changes in organelle architecture but also in the secretory pathways that produce SASP. Our surprising finding that LFU treatment of non-senescent cells induces the secretion of growth stimulatory factor(s) indicates that the mechanical effects of LFU may be part of a larger network of functions that support systemic responses to physical exercise. For example, exercise stimulates the secretion of myokines that benefit brain function and quality of life.^77^ We suggest that the effects of LFU may mimic many of the effects of exercise at a cellular level with the added benefit that LFU can penetrate the human body to reach internal organs.

The activation of growth of replicative senescent cells by non-invasive LFU has important implications for the *in vitro* expansion of normal cells to aid in autologous repair procedures, and it can augment the effects of nicotinamide on replicative senescence.^78^ Much more research is needed to understand the extent of the expansion that is possible. However, expansion does not involve apparent damage to the cells or major modifications of their phenotype. This indicates that LFU-induced reversal of senescence can have significant benefits in enabling the continued growth of normal cells beyond current limits.

As ultrasound has been approved for human exposure at power levels ten-to hundred-fold higher than the optimal levels used in this study, we suggest that it is practical to develop ultrasound-based therapies that could inhibit (or reverse) the increase in senescent cells in tissues with aging and thereby inhibit the onset of many age-related maladies. This non-invasive procedure has advantages over senolytics and exercise in that it is not tissue selective. In addition, LFU is non-invasive and will not directly affect biochemical or molecular biological treatments. Most importantly, these results show that mechanical treatments can augment or replace biochemical treatments to produce desired reversal of senescence and they are consistent with known effects of exercise on senescence and quality of life with aging.

## Acknowledgments

We want to acknowledge the experimental help of Adam Baker. We greatly appreciated the advice in editing this manuscript provided by Drs. Linda Kenney and Randy Levinson. Felix Margadant and Simon Powell provided support in fabrication of ultrasound devices and their maintenance.

## Funding

UTMB Biochemistry and Molecular Biology Department startup funds (MS)

CPRIT Foundation grant RR180025 (MS)

Welch Foundation professorship (MS)

Claude D. Pepper Older Americans Independence Center Pilot Project Grant (MS and BR)

NSF grant 1933321 (Co-I MS)

## Author contributions

Conceptualization: MS, SK, BR

Methodology: SK, RM, SP, FM

Writing: MS, SK, BR,

Experimentation: SK, RM, BB

## Competing interests

Authors (MS, SK, FM, and RM) are co-authors of patents related to these studies and MS and FM have financial interests in a company, Mechanobiologics, Inc that is planning to market LFU devices suitable for senescent cell rejuvenation.

## Data and materials availability

All data are available in the main text or the supplementary materials.

## Supplementary Materials

### Materials and Methods

#### Cell lines and cell culture

Human Foreskin Fibroblasts (HFFs) and Bone marrow-derived mesenchymal stem cells (MSCs)were purchased from the ATCC. African monkey kidney-derived Vero cells were obtained from the M. Garcia-Blanco lab as a gift. All these cell lines were cultured as per manufacturer’s protocol. Vero cells and HFFs were in growth medium containing Dulbecco’s Modified Eagle’s Medium (DMEM) 10% fetal bovine serum (FBS; Gibco) and 1% Penicillin/Streptomycin. Human MSCs were cultured in MSCs-approved medium (ATCC) and expanded as per the supplier’s protocol. Culture medium was changed every 48 h unless otherwise stated. Cells plated at 20-40% confluency were maintained in an incubator at 37^0^C and 5% CO^2^. Cells were passaged every 48-72 h using Trypsin/EDTA (Gibco). Cells were counted manually using a hemocytometer and ImageJ in at least three independent replicates unless otherwise stated. Minimally 5 fields were counted from each replicate.

#### MSC differentiation assays

LFU-treated and untreated P18 MSCs were cultured in a 12-well culture plate in growth medium for 24 h. Then, growth medium was replaced by adipogenic (Invitrogen) or osteogenic (Invitrogen) differentiation media as per the manufacturer’s protocol. Adipocytes were assayed after 12 days by Oil Red O (Sigma Aldrich) staining and osteocytes were assayed using an Alizarin red S dye (Sigma Aldrich) solution. Images were acquired using a 10x evos RGB objective.

#### Senescence induction and quantification

Vero cells were treated with various stressors, including 200 μM H_2_O_2_, 4 mM of sodium butyrate (SB), 25 mM bleomycin sulphate (BS) or 200 nM doxorubicin and incubated for 36-48 h. After washing with PBS and then adding fresh medium, cells were incubated for 4 days to confirm the growth arrest of senescent cells (*59*). HFFs were serially passaged until P15 as replication of these cells was dramatically reduced by P15-17. We used four criteria to determine if cells were senescent; (1) Cell cycle arrest by determining the growth rate, (2) Increase in cell spread area, (3) Development of a senescence-associated secretory phonotype (SASP) in culture medium and (4) β-galactosidase staining. We captured images of cells with an Evos microscope at 10X magnification after LFU treatment and 48 h post treatment. To measure growth by the increase in cell number, 15 random images were captured, then the average number of cells were determined, which was divided by the area of one frame to get the cell density (cells/cm^2^). Then this seeding density was multiplied by the total area of the dish or well to obtain the total number of cells after LFU treatment and after 48 h of incubation. The total number of cells at 48 h was divided by the total number of cells just after LFU treatment to determine the growth rate. If the ratio was one, there was no growth.

Senescence was detected by the β-galactosidase senescence staining kit as per the manufacturer’s protocol. Briefly, sub-confluent senescent cells were stained by the SA-β-galactosidase staining solution and incubated overnight at 37^0^C. The β-galactosidase-stained cells appeared blue and were considered senescent cells. The percentage of β-galactosidase-positive cells was determined by counting the number of blue cells and dividing by the total number of cells. Cell spread area was determined by capturing the images of cells with a 10X objective using an Evos microscope. Then, we used ImageJ software to calculate spread area by manually encircling the cell periphery of each cell. We used a minimum of 150 cells for the analysis. To determine the SASP activity, we cultured the senescent cells for 3-4 days and then supernatant was collected from each dish. This supernatant was used to culture normal cells. Development of a senescence phenotype by the normal cells in the supernatant medium confirmed that senescent cells 55 were secreting SASP.

#### LFU treatment of cells

Prior to LFU treatment, the plates containing senescent Vero cells or late passage HFF cells were wrapped with parafilm to avoid contamination and water influx into the plate. The samples were placed on the plastic mesh, which was mounted on the water tank with an ultrasound transducer. Water in the tank was degassed and heated to 35^0^C. The distance between the sample and transducer was approximately 9-10 cm. We also ensured that there were no air-bubbles or air-water interfaces between the water and the sample. Output power of the transducer was measured at the plate location by a calibrated needle hydrophone (ONDA MCT-2000). Cells were treated with pressure pulses of intermediate power and low frequency for 30 mins. Cells were treated with a 50% on-off duty cycle. After LFU treatment, cell plates were returned to the incubator for 48 h to determine the growth of senescent cells.

#### Reversal of senescence

Firstly, we induced senescence in Vero cells using sodium butyrate, then we confirmed senescence using growth arrest and β-galactosidase staining after four days of incubation which normally eliminated quiescent cells (*59*). The senescent cells were treated by LFU with optimized parameters (power, frequency, and duty cycle) and incubated for 48 h to measure the growth and morphology of the cells before trypsinization (passage P0). These cells were then trypsinized, reseeded and incubated for 48 h for the P1 passage. This process was repeated to a P3 passage. Growth in number, morphology, β-galactosidase and EdU incorporation were measured and all indicated that LFU reversed the senescence. Control cells without LFU treatment remained senescent. Typically, by P3, the senescent cells exhibited the phenotype of normal proliferating cells.

Passage 15-24 HFFs were treated with LFU at the optimized frequency and power. Cell proliferation was determined by counting the number of cells at the time of seeding and 48 h post LFU treatment. LFU-treated HFFs showed a higher growth rate than the untreated HFF cells. In the case of control P24 HFFs, they were treated with LFU and incubated for 96 h prior to trypsinization, reseeding and incubation for 48 h. Proliferation and morphology were measured after 48 h of incubation. P24, LFU-treated HFF cells became smaller in size, and they also showed dramatically greater proliferation than the untreated P24 HFFs.

#### Soft surface preparation

PDMS of 5 kPa elastic modulus was prepared by mixing the Sylgard 184 silicone elastomer kit at an Elastomer/curing ratio in 75:1. The combination was mixed, degassed and spin coated at 4000 rpm for 20 seconds on 27 mm ibidi glass bottom dishes. PDMS coated dishes were incubated at 65°C overnight. Dishes were cleaned, activated by oxygen plasma treatment, then coated with 25 μg/ml Fibronectin (Sigma Aldrich) and kept overnight at 4°C. Coated dishes were washed with PBS before plating the cells.

#### Microtubule analysis

Senescent cells were treated with tubulin tracker deep red (Invitrogen) as per manufacturer’s protocol. Briefly, cells were incubated in tubulin tracker at 1000:1 for 30 min and medium was changed before confocal imaging with a 60x oil objective. The cells were treated with LFU and imaged again at the same confocal microscope settings. Length of myotubules was determined as described previously (^52^.

#### Pharmacological drug treatment

The following inhibitors were used in the study: Nocodazole (1 μM), Cytochalasin D (100 nM), Blebbistatin (10 μM), Y27632 (1 μM), EX-527 (10 μM), GsMTx (1 μM) and Rapamycin (1 μM). Senescent cells were incubated in Resveratrol (100 μM) for 24 h to activate Sirtuin1 activity. L-Leucine (10 μM) and Rapamycin (200 nM) were used to activate and inhibit phosphorylated mTOR during the 30 min LFU treatments. To assess LFU effect on TRPV1 activation or inhibition, we used Ruthenium Red (10 μM) and Capsaicin (10 μM) during the LFU treatments. For these treatments all inhibitors were purchased from Sigma Aldrich and prepared in DMSO and milli-Q water as per the manufacturer’s protocol. Cells were incubated in inhibitors overnight after 6-8 h cells seeding prior to LFU.

#### Senescence assay

Senescence was detected by the senescence-associated β-galactosidase staining (Sigma Aldrich) as per manufacturer’s protocol. Briefly, senescent, non-senescent and LFU-treated cells were seeded in 27 mm ibidi glass bottom dish and incubated for 48 h. Then, these cells were fixed and stained per the manufacturer’s protocol. After adding staining solution into the dish, cells were incubated overnight at 37°C in absence of CO_2_. A minimum of 100 cells were counted manually for each condition of the analysis. 10–15 random images were captured for each condition for analysis of β-galactosidase-positive cells, which were counted manually.

#### EdU and Annexin V staining

Cells were incubated with 10 mM EdU reagent for 24 h. Cells were then fixed, permeabilized and blocked according to the manufacturer’s protocol (Click-iT EdU Alexa Fluor 555 imaging kit, Life Technologies). Hoechst was used to stain nuclei at a 5000:1 ratio. Images were captured with a 10X objective. The number of red puncta over the total number of blue nuclei gave the percentage of EdU-positive cells. At least 200-300 cells were counted manually using ImageJ for each analysis. Apoptosis of senescent cells was determined using Annexin V Alexa fluorophore 488 staining solution. Senescent cells were treated with LFU and after 30 m, they were stained for Annexin V according to the manufacturer’s protocol and images were taken with an Evos microscope at 10X. Senescent cells were treated with 0.5 mM H_2_O_2_ as a positive control.

#### Mitochondrial morphology

Ultrasound-treated cells were incubated with Mitotracker (Invitrogen) at 250 μM at 37°C for 30 min. Then, images were captured in by a confocal microscope for quantification (15 randomized fields per sample). Mitochondrial velocity was determined from the time lapse image by an Olympus microscope at 60X. Images were captured after every 5 seconds for 10 minutes. Mitochondrial mobility was determined by ImageJ software with manual tracking. 10-15 mitochondria were traced manually.

#### Immunofluorescence staining

Cells were fixed with 4% paraformaldehyde for 15 minutes, permeabilized with 0.5% Triton-X for 5 min and blocked with 3% Bovine serum albumin (BSA) for 1 h at room temperature and then incubated in primary antibody overnight at 4^0^C. The following primary antibodies were used: rabbit polyclonal p21 antibody (cell signaling) (1:400), Mouse monoclonal p16 antibody (1:300) (Abcam), Mouse H3k9me3 (1:500) (ThermoFischer), Mouse p53 (1:200:) (SantaCruz Bio), Rabbit Gamma H2XA (1:200:) (Cell Signaling), Mouse monoclonal SIRT1 antibody (1:300:) (Sigma), p-mTOR, Ser 2448 (Cell-signaling D9C2) (1:200:), Rabbit Anti-YAP (ab52771) (1:300:), α-Tubulin (ab7291) (1:500:). After washing with PBS, cells were incubated with goat anti-mouse 488 (1: 1000) (Sigma) and Donkey anti-rabbit 555 (1: 1000) (Invitrogen) for 2 h. Nuclei were stained with Hoechst at a 1: 5000 dilution. Images were acquired by spinning-disk confocal microscope. Mitotracker green (Invitrogen) at 100 nM, Lysosome tracker deep red (Invitrogen) at 50 nM, and Tubulin tracker deep red 1: 1000: were used as fluorescent tags.

Sodium butyrate-treated HFF cells were incubated in MitoSOX Red (Invitrogen) and ROS probe (Dojindo’s ROS Assay Kit) as per manufacturer’s protocol. Briefly, cells were treated with 5 mM of MitoSOX Red and 1:1000 ROS detection dye in HBSS for 30 mins. Then, cells were washed twice with HBSS and treated with LFU and Cells were imaged at Olympus microscope using Ex/EM 490/520 and EX/EM 396/610 in controlled environment at 5% CO_2_ and 37^0^C.

#### Cytokine profiling by multiplex cytokine assay

To analyze the level of chemokines, cytokines and immunoregulatory proteins in the supernatant of LFU-treated late passage and P3 HFFs, we used Bio flex cytokines 27-flex kit assay (Bio-Rad). This multiplex immunoassay assay contained fluorescence nanoparticles conjugated with a specific antibody and was used according to the manufacturer’s protocol. Briefly, supernatant collected from the late passage HFFs and P3 cells treated with or without LFU, were diluted 1:4 and incubated for 2 h at room temperature in shaking. Then, wells containing samples were washed, and incubated with biotinylated antibody for 1 h at room temperature, then with streptavidin-phycoerythrin for another 30 min. A calibration curve was established using the standard given with the assay kit. Bioplex-200 plate reader (UTMB Galveston, Texas core facility) was used to assess the cytokine level. The concentrations of molecules were determined using the standard curve provided by the manufacturer.

#### Analysis of autophagic flux

To analyze LFU-induced autophagy in late passage HFFs and in LFU treated HFFs, we used Prema autophagy Tandem sensor RFP-GFP-LC3 kit (Invitrogen). Tandem sensor has acid sensitive GFP and acid insensitive RFP. In active autophagy, fusion of lysosomes with autophagosomes increases acidity, which quenches the GFP. Cells were cultured in 35 mm glass bottom dishes for 24 h before LFU treatment. Cells were transduced with autophagy tandem sensor according to the manufacturer’s instructions. Briefly, 40 particles per cells were used for transduction, and incubated overnight in culture medium. Chloroquine diphosphate (100 μM) was used as an autophagy blocker overnight. Cells were visualized and images were captured at 100x on a spinning-disk confocal microscope (Olympus). Autophagy was determined as the ratio of intensities of GFP and RFP.

#### Telomere length measurement

The length of telomere was determined using Absolute Human Telomere Length Quantification qPCR Assay Kit (ScienCell AHTQL-8918) according to the manufacturer’s protocol. Total genomic DNA was extracted from Human foreskin fibroblasts of various conditions including early passage (P2), late passage (P18) cells, LFU-treated and untreated control cells using Invitrogen Pure Link Genomic DNA kit (K182001). 5 ng of genomic DNA was used as a template in PCR. Every experimental genomic DNA was processed four times. The reference DNA was analyzed in two triplicates. Telomere length was calculated according to the manufacturer’s protocol.

#### DNA methylation assay

The protocol for DNA methylation was performed as reported previously.^79^ Cells were incubated in 20 µM CldU for 16 h. After trypsinization, cells were harvested and resuspended in ice-cold PBS. A 2 µl droplet containing 200-300 cells was diluted in 8 µl of Lysis buffer (0.5% SDS, 50 mM EDTA, 200 mM Tris-HCl, pH 7.4). A 10 µl droplet containing cells diluted in Lysis buffer was poured onto the silane prepared slide (Sigma Aldrich), tilted, and allowed to flow by gravity. The slide was then fixed in methanol/ acetic acid (3:1) for 25-30 min and rinsed with PBS thrice. Slides were then denatured in 2.5 M HCl for 1 h; neutralized in 0.4 M Tris-HCl for 5 min and blocked in 5% BSA buffer for 1 h. Slides were stained with anti-BrdU (1:200) and 5mc (1:200) antibodies overnight at 4°C. Slides were stained with Alexa flour 488 goat anti-rat (1:1000) and Q-dot 655 goat anti-mouse (1:2000) for 1 h for BrdU and 5mc antibody respectively.

#### RNA extraction and sequencing

RNA was extracted from early passage HFFs (P2, P3, and P4) and rejuvenated senescent cells (P18, P19, and P20) using Qiagen RNeasy Mini Kit. (Germantown, MD). The UTMB Next Generation Sequence (NGS) core laboratory assessed RNA concentrations and quality using a Nanodrop ND-1000 spectrophotometer (Thermofisher, Waltham) and an Agilent Bioanalyzer 2100 (Agilent Technologies, Santa Clara, CA). PolyA+ RNA was purified from ∼100 ng of total RNA and sequencing libraries were prepared with the NEBNext Ultra II RNA library kit (New England Biolabs) following the manufacturer’s protocol. Libraries were pooled and sequenced on an Illumina NextSeq 550 High Output flow-cell with a single-end 75 base protocol. Reads were mapped to the human GRCh38 reference genome with STAR version 2.7.10a with the parameters recommended for the ENCODE consortium. Reads mapping to genes were quantified with the STAR –quantMode.^80^ GeneCounts option using the Gencode v41^81^ primary assembly annotation file. Differential gene expression was estimated with the DESeq2 software package, version 1.38.3, following the vignette provided with the package.^80^ The complete RNA-seq data will be posted online.

#### Statistical analysis

All experiments data reported were obtained from the minimum of three samples or pre group otherwise mentioned in the figure legends. Data is represented as the mean ± standard deviation and Statistical analysis was performed using GraphPad prism 10.0. Differences in the two group is determined using the two-tailed paired, unpaired Student’s *t*-test, Mann Whitney tests, and Kruskal Wallis, and One way ANOVA test followed by post hoc Dunn’s test used for the multiple groups. Statistical significance was analyzed and reported in the figures and figure legends *, *P* values < 0.05, **, *P* values < 0.002, ***, *P* values < 0.001, ****, *P* values < 0.0001 and non-significant (ns) *P*-value > 0.05.

---

**Figure S1.**
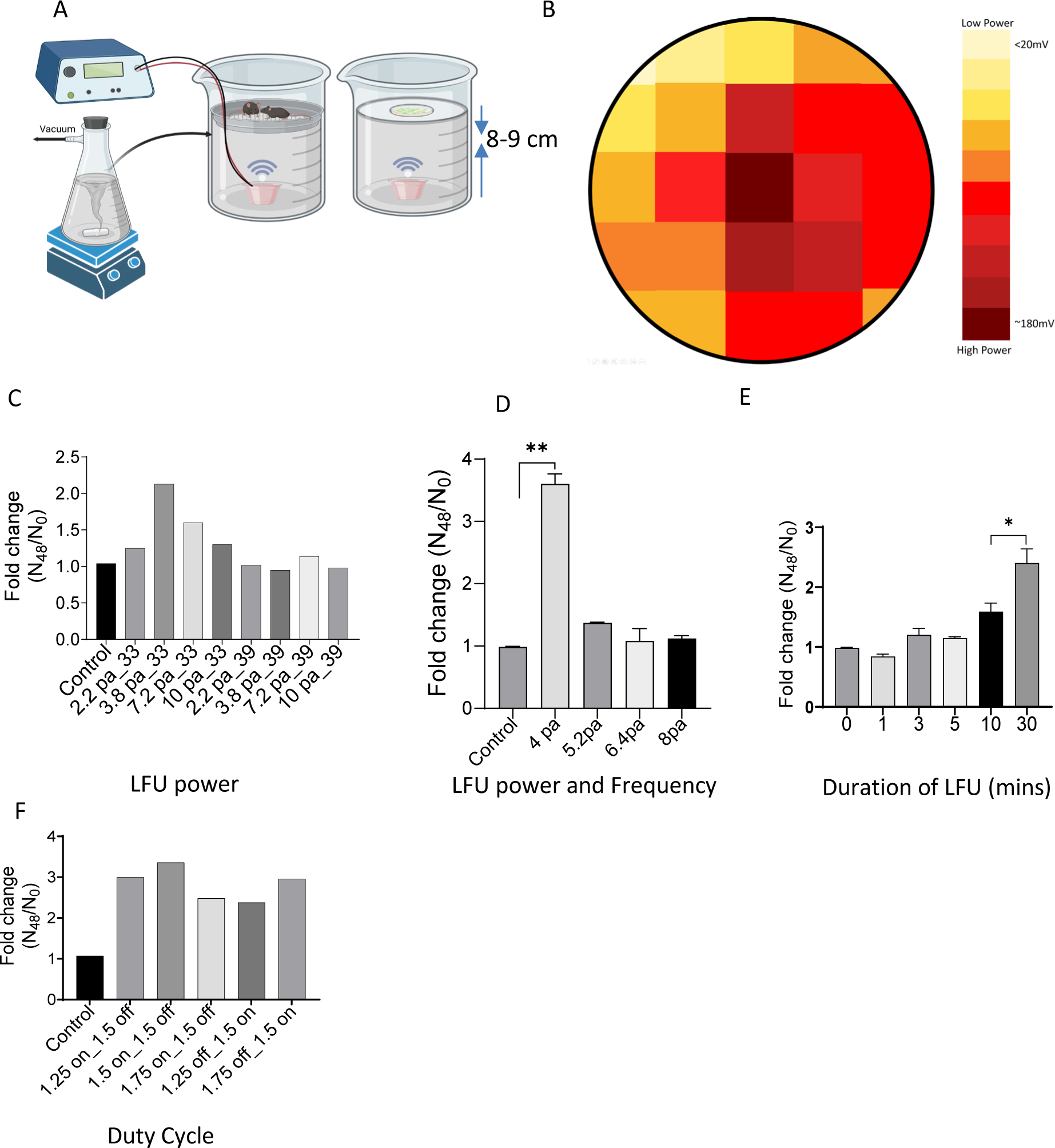
Low Frequency Apparatus and Optimization Experiments. (A) Schematic of LFU treatment setup for cells and mice. (B), Heat map representing the distribution of LFU output power at the location of treatment, darkest red shows the maximum output power measured by the hydrophone and the yellow color shows the minimum output power. (C), Bar graph showing the growth of Sodium butyrate senescent Vero cells (SCs) in 48h hr after LFU treatments with various power levels at 33 & 39 kHz. Data is represented as the mean of three replicates. (D), Bar graph representing the growth of SCs in 48 h after 33 kHz LFU treatment at the designated power level. Data is represented as the mean ± s.d. of three independent experiments. ** p-value < 0.01 by unpaired two tailed student t-test. (E), Growth was determined as a function of the treatment duration. Control represents no LFU treatment. Results plotted are the mean of three independent experiments ± s.d. *p value <0.05 by unpaired two tailed student t-test. (F), Effect of duty cycle on growth, where control is without LFU. The power level was 4 Pa at 33 kHz for 30’. Results plotted are mean of 3 replicates.

**Figure S2.**
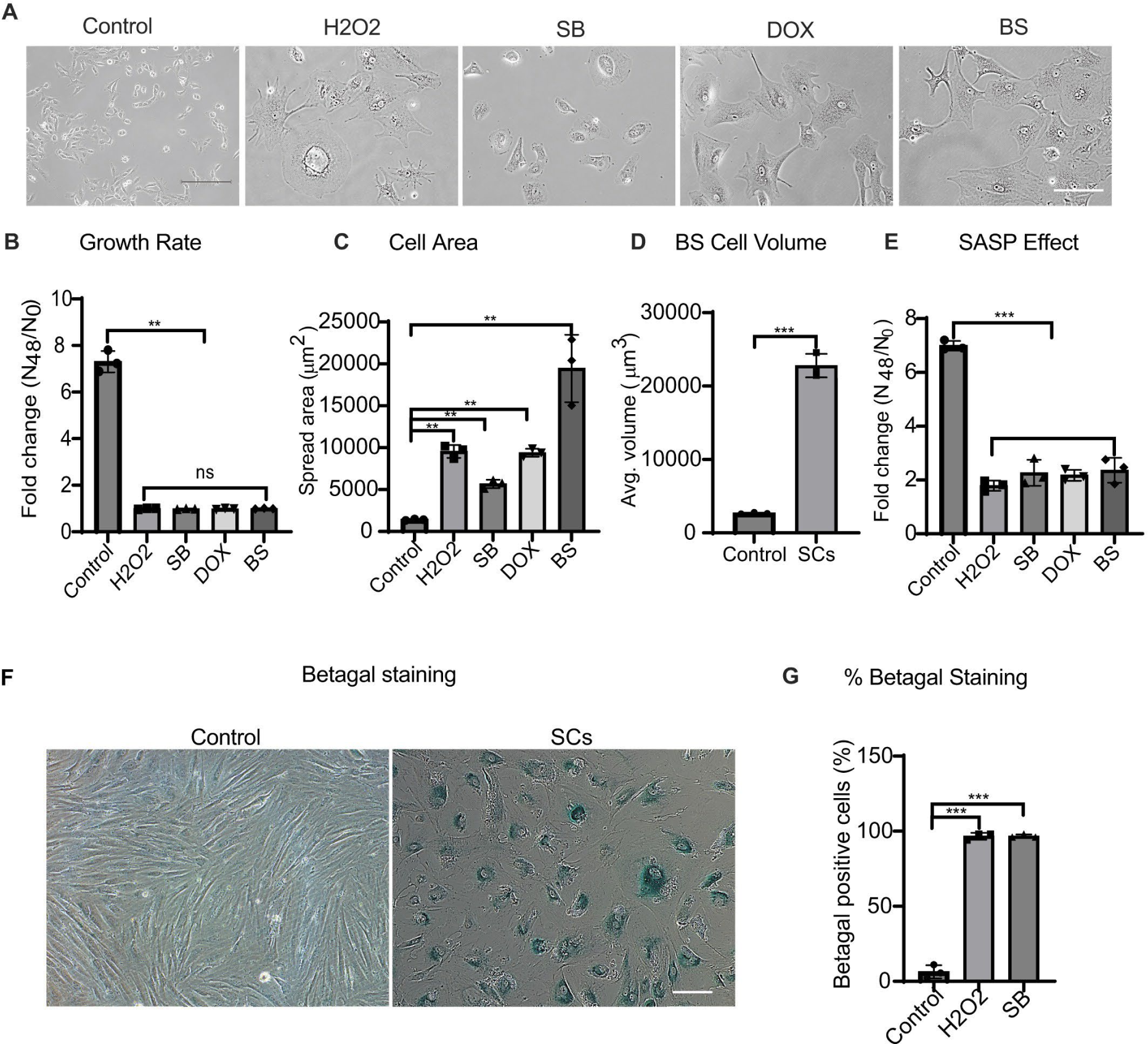
Characterization of Senescent Cells. (A) Brightfield images of control Vero cells and senescent cells induced by H_2_O_2_, sodium butyrate (SB), Doxorubicin (Dox), and Bleomycin Sulphate (BS). Scale bar=300 µm. (B) Quantification of proliferation shows no senescent growth after a 48 h incubation. (C)Senescent cells become enlargedcompared to the normal control cells. (D) Quantification of avg. cell volume of bleomycin sulfate treated cells compared to control cells. (E) Supernatant collected from the senescent cells after 24 hour incubation was used to study the growth of normal proliferating Vero cells. Culture medium was used as a positive control. Result are the mean of three experiments ± SD. (F) SA-b-galactosidase staining of control (proliferating) and BS treated senescent cells. Scale bar= 300 µm. (G) Level of b-galactosidase senescence marker in H_2_O_2_ and SB induced SCs. Scale bar=300µm. *P<0.05, **P<0.002, *** P<0.0001;data in (B)-(E) and H are mean ± SD. Minimum of 150 cells were analyzed for spread area, cell volume and b-galactosidase from three independent experiments.

**Figure S3.**
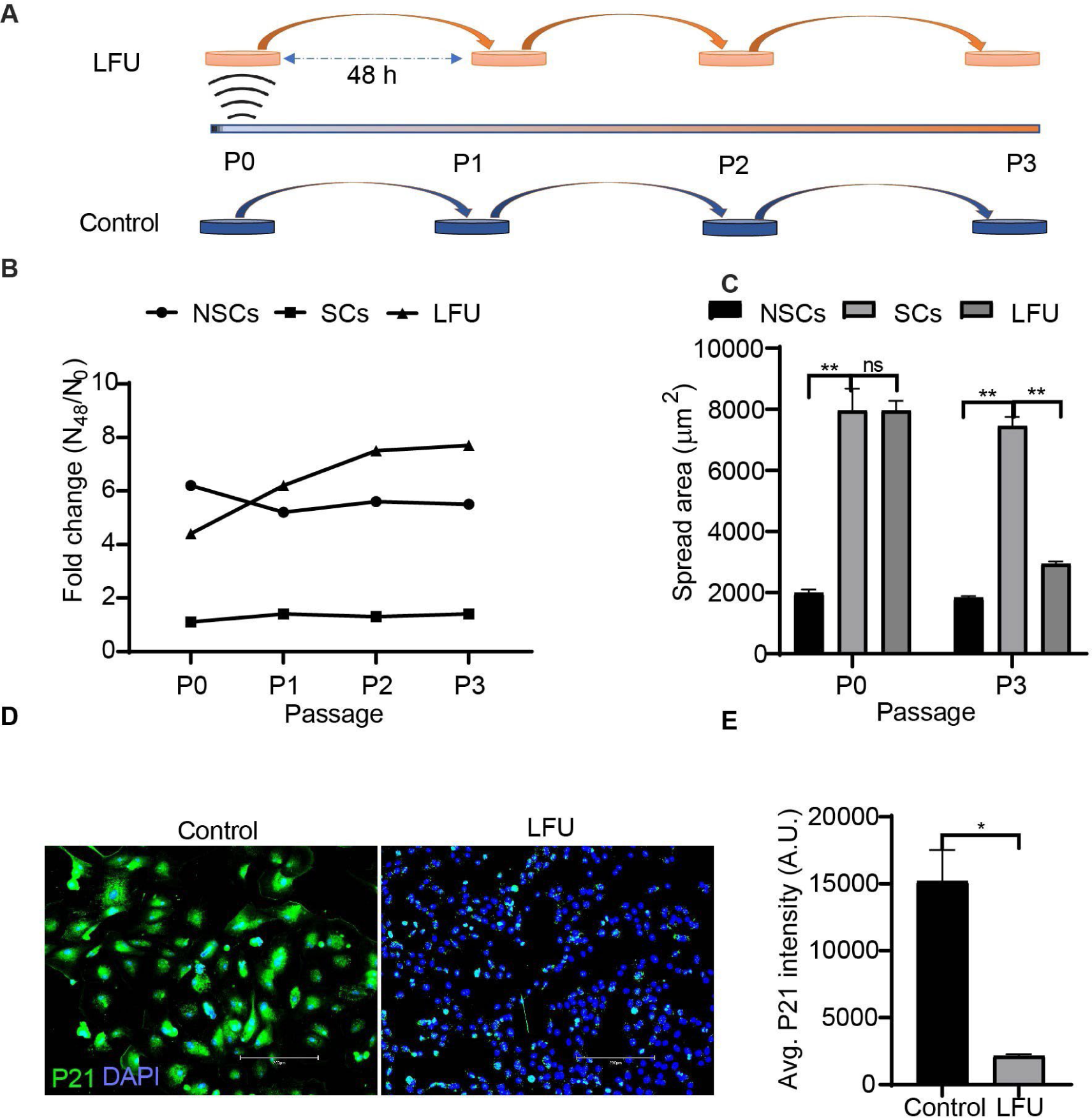
Low Frequency Ultrasound (LFU) Reverses Cell Senescence. (A) Schematic illustration showing experiment design of LFU induced reversal of senescence after Sodium Butyrate (SB) treatment for 48 hour followed by incubation for 4 days. Cells were treated with LFU for 30’ and then passaged every two days. (B) Growth of SB treated senescent Vero cells after LFU treatment and passage every 48 h for 8-10 days. NSCs were the non senescent Vero cells. Graph shows growth of normal Veros (NSC), BS senescent (SC) and LFU treated (LFU) senescent cells as fold change over 48 h for passages from P0 to P3 every 48 h. (C) Cell area of LFU treated senescent cells (LFU) is largely restored to normal by P3. (D) Representative anti-p21 immunofluorescence images of senescent P3 control and LFU treated senescent P3 cells. Scale bar= 300 µm. (E) Quantification of fluorescence intensity of p21 stained senescent and LFU treated senescent P3 cells shown as mean ± SD, for >200 cells in each condition. All graphs were plotted by mean ± SD and p values ns p>0.05, *P<0.05, ** P<0.002. Minimum 200 cells were analyzed from three independent experiments in each condition.

**Figure S4:**
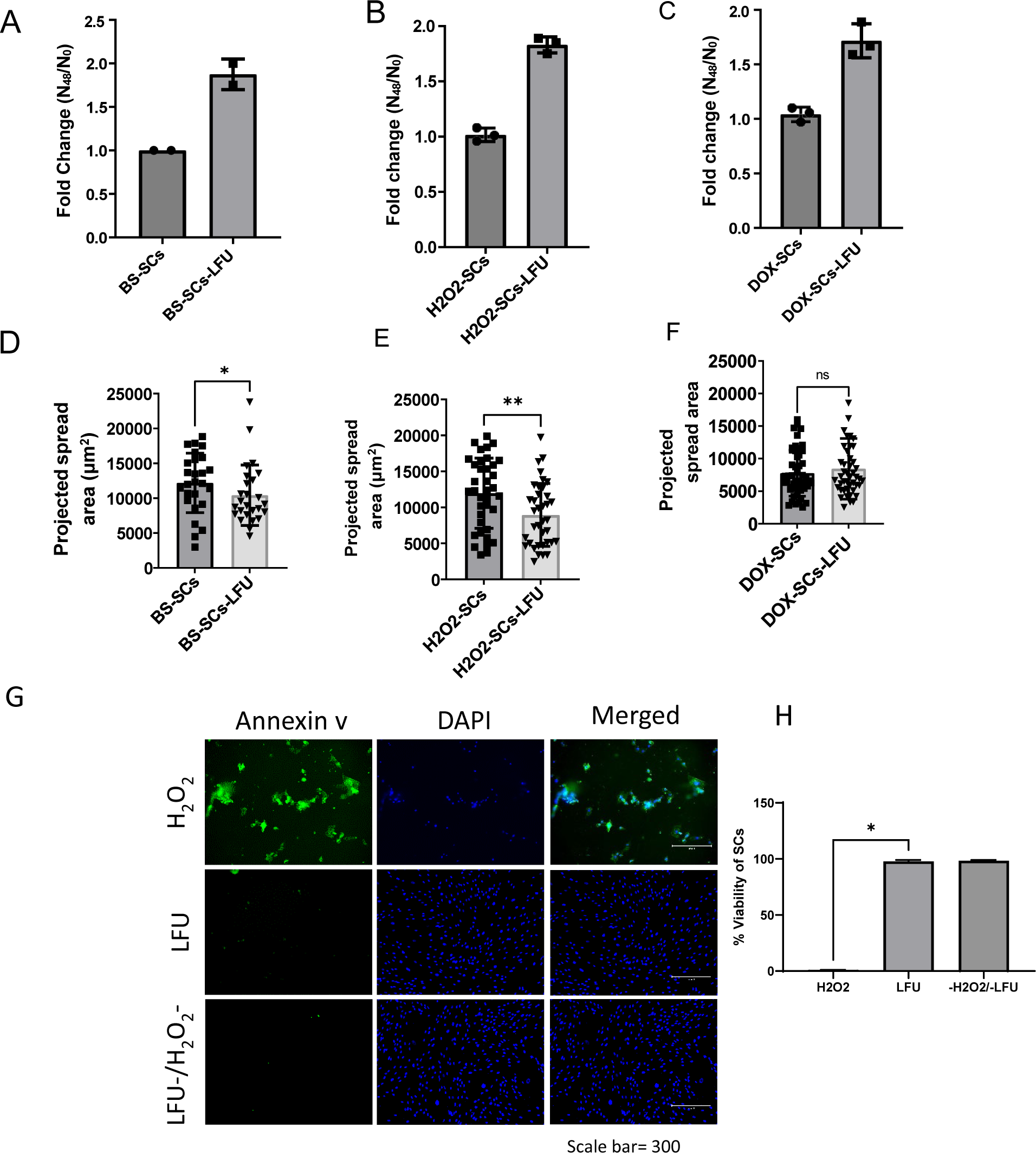
LFU treatment activates growth and reduces cell size without apoptosis. (A, B,C) Growth and Area of Bleomycin Sulphate (BS) (50 μM) cells treated w/wo LFU after 48 h. (D, E, F) Growth and Area of H_2_O_2_ (200 mM) cells treated w/wo LFU after 48 h. (E, H) Growth and Area of Doxorubicin (200 nM) cells treated w/wo LFU after 48 h. (G) Annexin v and DAPI stained vero cells treated with nothing, 10 mM or 200 mM H_2_O_2_ plus LFU. (H) % viable cells after LFU with nothing, 10 mM or 200 mM H_2_O_2_. Results are plotted as mean of three replicates and ± s.d. At least 35-50 random cells were analyzed from the three replicates. Non-parametric Mann Whitney test was used to determine the statistical difference between the two groups. * p value <0.05. ** p value<0.001 and ns p value >0.05.

**Figure S5:**
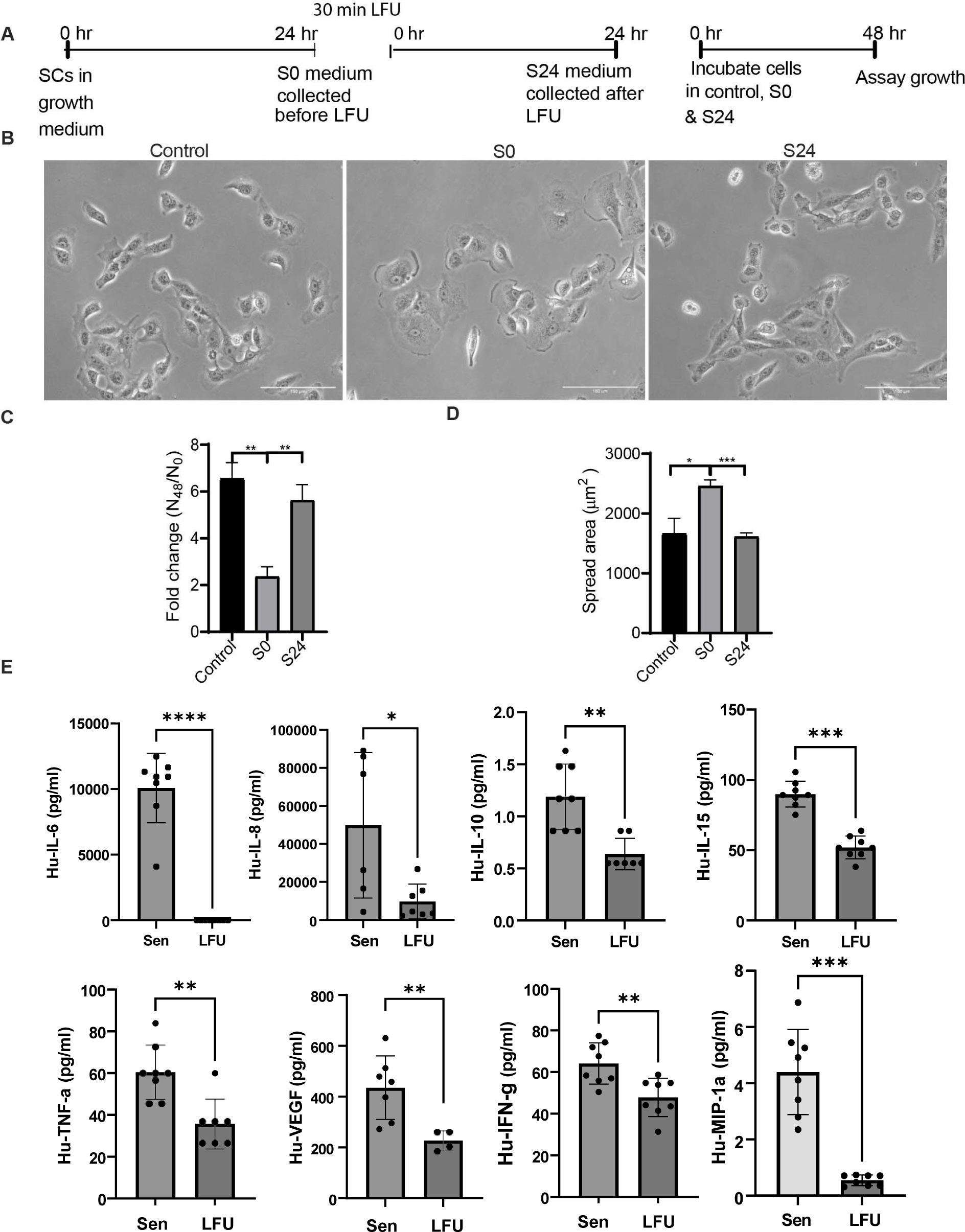
LFU Treatment Decreases SASP Secretions. (A) Schematic illustration of the experiment where BS senescent cells were cultured in growth medium for 24 hr and then they were treated with low frequency ultrasound (LFU) for 30 min. Supernatant was collected after the LFU treatment (S0) and cells were incubated for another 24 hour in fresh medium before supernatant was collected (S24). To check the effect of LFU treatment, supernatants S0 and S24 were used to check the growth of non-senescent HFF cells. (B) Representative brightfield images of normal vero cells after 48 hours of incubation in control growth, S0, or S24 medium. (C) Quantification of normal vero cell numbers after 48 in control growth, S0 or S24 medium. (D) cell areas after 48 in control growth, S0 or S24 medium. (E) Chemokines and cytokines in supernatants from untreated and LFU treated late passage HFF (P19) cells after 24 hours incubation were measured using Multiplex assay. Results are plotted as mean± s.d., n= 6 replicates, ns not significance, * p value <0.05 ** p value <0.01, *** p value 0.0001, and **** p value 0.00001 using Mann Whitney Test.

**Figure S6:**
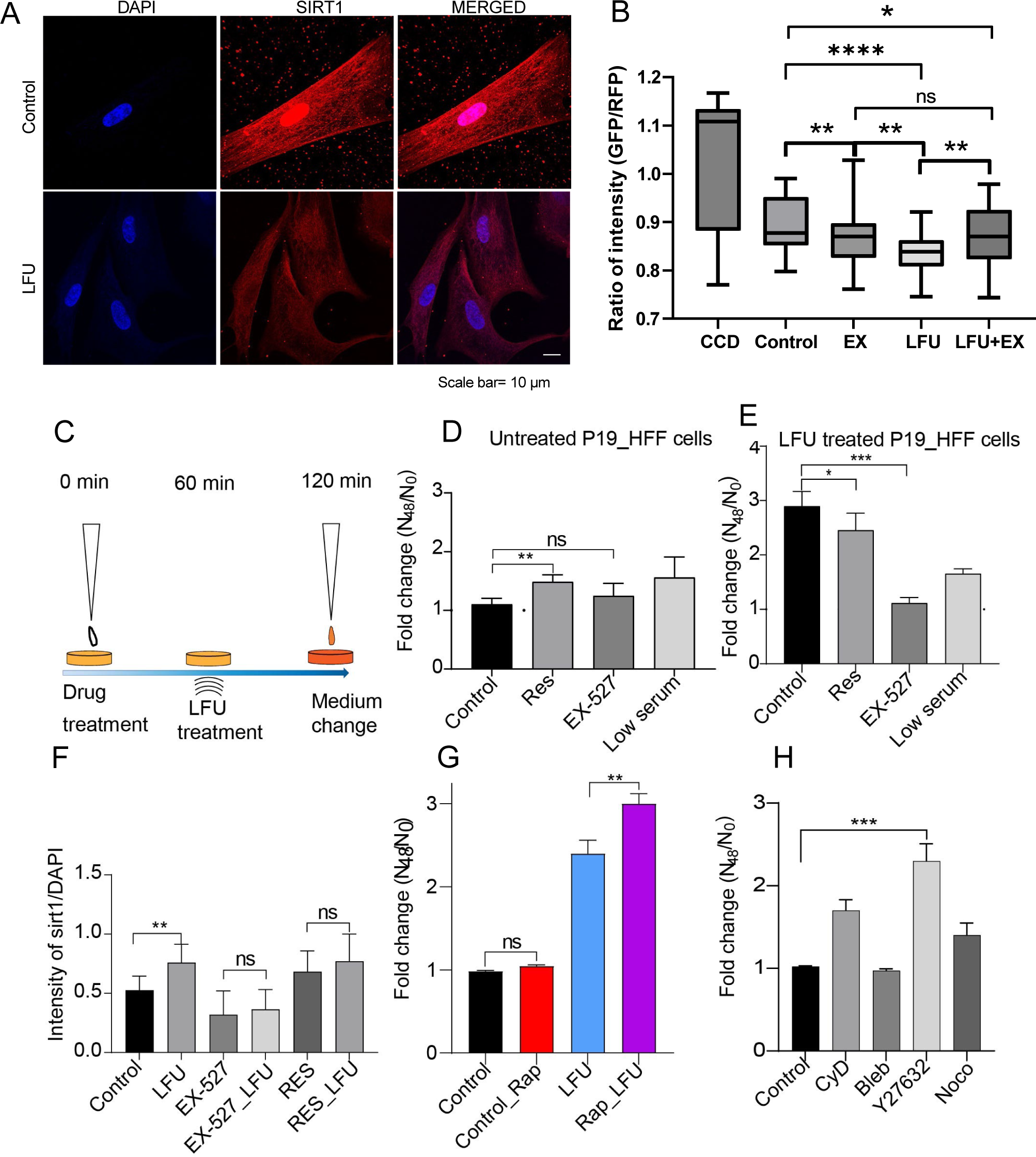
Sirtuin 1 Distribution and Function in Rejuvenation of Growth and Autophagy. (A) Distribution of Sirtuin 1 antibody and DAPI in senescent and LFU rejuvenated cells. (B) Ratio of GFP/RFP in BS cells transfected with GFP-LC3-RFP after treatment with nothing (Control), 10 mM Chloroquine diphosphate (CCD), 10 mM of EX-527 (EX), LFU for 30’, or EX-527 plus LFU (LFU-EX). (C) Schematic of Resveratrol and EX-527 treatment, followed by Ultrasound treatment and medium change. (D) Growth after 48 h of untreated control senescent P19_HFF cells in presence of 10 mM resveratrol (Res), 10 mM of EX-527 (EX-527), or low serum (1% serum). (E) growth after 48 h of LFU treated P19_HFF cells in the presence of the same drugs. (F) quantification of Sirtuin 1 expression by intensity ratio of Sirtuin 1 antibody fluorescence to DAPI fluorescence after no treatment (Control), LFU treatment for 30’ (LFU), 10 mM of EX-527 treatment (EX-527), EX-527 and LFU treatment (EX-527_LFU), Resveratrol treatment (RES), and Resveratrol plus LFU treatment. (G) Growth rate after 48 h of P19 HFF (Control), treated with Rapamycin (1 mM) (Control_RAP), treated with LFU (LFU) or treated with Rapamycin plus LFU (RAP_LFU) shows significantly increased growth with rapamycin and LFU treated SCs compared to with and without rapamycin controls. (H) Growth of P19 HFF SCs in presence of cytoskeleton inhibitors including Cytochalasin D (10 mM), Blebbistatin (10 mM), Y27632 (1 mM), and Nocodazole (1 mM).

**Figure S7.**
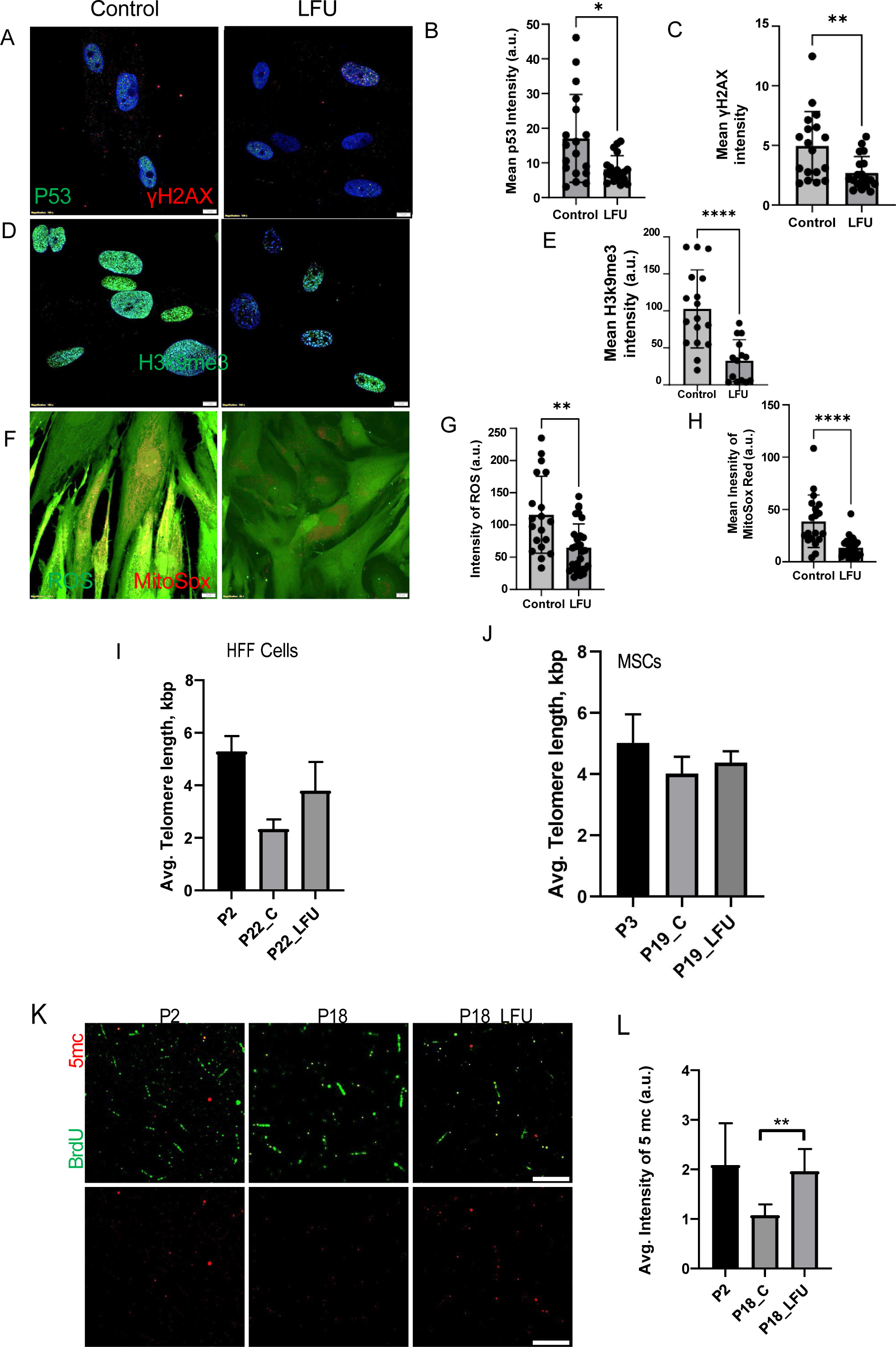
Senescence Markers Are Reduced, Telomers and 5mc Are increased after LFU Treatment. (A) Representative images of p53 (green) and γH2AX foci (red) stained cells. Sodium Butyrate senescent Vero cells were treated two times with or without LFU and then immunostained with P53 and γH2AX antibodies. Staining was done 24 hours after LFU treatment (2^nd^ treatment that was 1 day after first). (B) Quantification of p53 (green) intensity from the three replicates. (C) Quantification of γH2AX mean intensity from three replicates. (D) Representative images of H3k9me3(green) stained cells Sodium Butyrate senescent Vero cells were treated two times with or without LFU and then stained with H3k9me3 antibody (one day separated first and second LFU treatments and staining). (E) Quantification of H3k9me3 (green) intensity from the three replicates using standard conditions. (F) Representative images of ROS (green) and MitoSOX (red) treated cells with live cell staining probe. Sodium Butyrate induced Vero cells were treated with or without LFU and images were captured after 24 h. (G) Intensity of ROS (green) staining and (H) MitoSOX (red) staining from the three replicate experiments. (I) Telomere length measurements of HFFs after passage 2 (P2) and passage 22 (P22) shows shortening but length increases with LFU (P22_LFU) n=4, N=1. Results are shown as mean ± SD. (J) In mesenchymal stem cells, the telomere length decreases slightly from passage 3 (P3) to passage 19 (P19_C) but is increased proportionally by LFU (P19_LFU), n=6 from one experiment. Results are shown as mean ± SD from the three independent experiments. (K) Fluorescence images of passage 2 (P2), untreated P18 (P18_C) and LFU treated P18 cells (P18_LFU) HFF cells with BrdU and 5 mc fluorescence. (L) Quantification of intensity of 5mc fluorescence. Scale bar =10 μm. Data is plotted as mean± s.d. from the three replicates. One data point represents one cell analyzed. Non-parametric Mann Whitney test was used to determine the statistical significance between two groups. * p value<0.05, ** p value <0.007, **** p value<0.0001, and ns p value >0.05.

**Figure S8.**
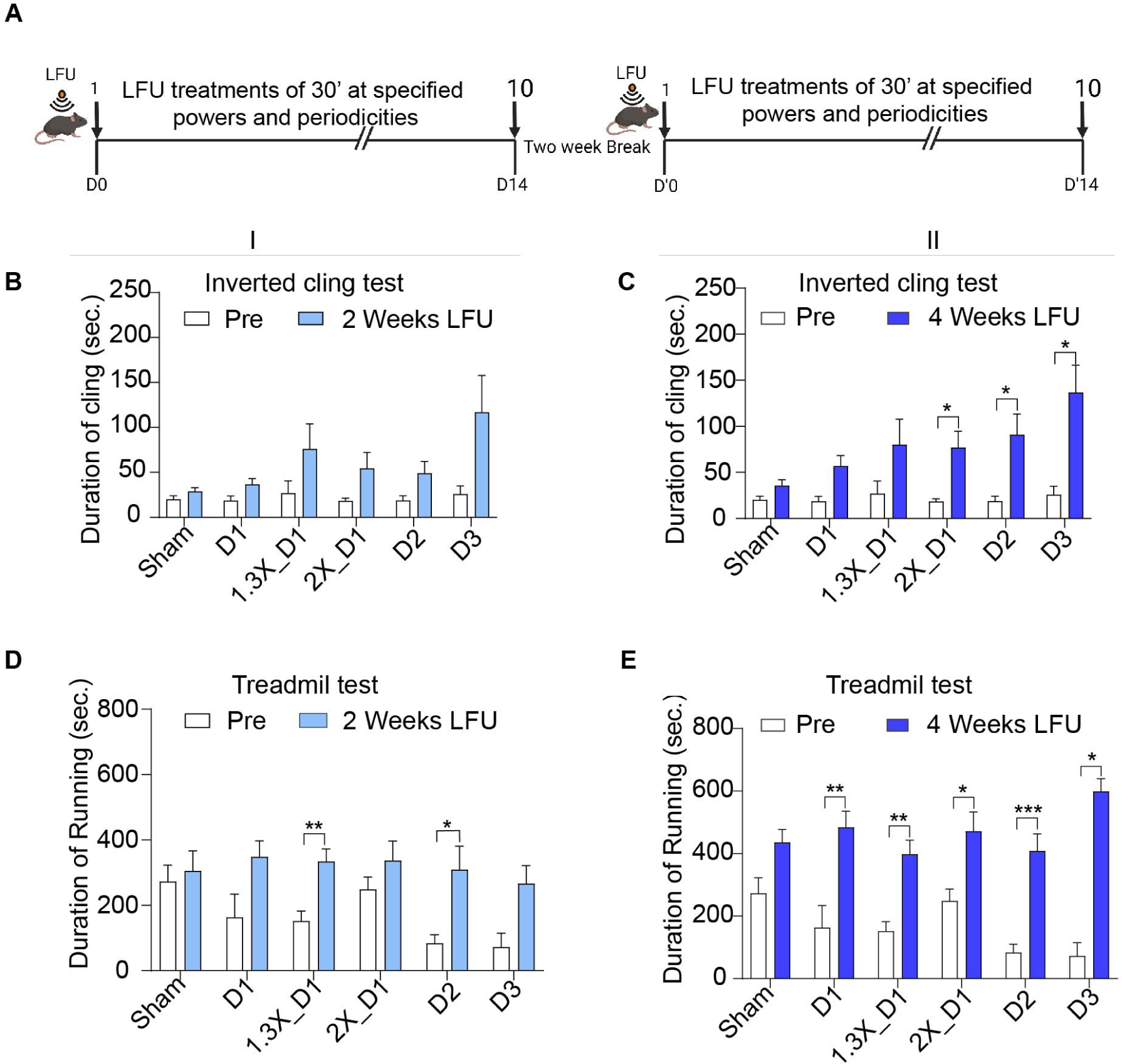
Effects of LFU dosage on performance of aged mice. (A) Schematic illustration of the treatment plan. Mice (20-24 months old mice (C57BL/6J strain)) were treated 30 min at 1X power every day (D1), every other day (D2) and every third day (D3), 1.3x power every day (1.3X_D1) or 2X power every day (2X_D1) for two weeks followed by a two week break and then two-weeks of treatment. Performance of treated mice was assessed as pre and post treatment with standard tests. Each study group contained five male and five female mice. (B) Results of inverted cling test after the first two weeks of LFU treatment. And (C) Inverted cling test data after four weeks of treatment with 2 weeks break. (D) Results of the treadmill test after the first two weeks of LFU treatment (see Materials and Methods mouse protocols). (E) Results of the treadmill test (see Materials and Methods mouse protocols) data after four weeks of treatment with two weeks break. Results are plotted as mean ± SEM. Student t-test was used to determine the statistical significance. Statistical significance was given in p-value. *P<0.05, **P<0.002, *** P<0.0001.

**Figure S9.**
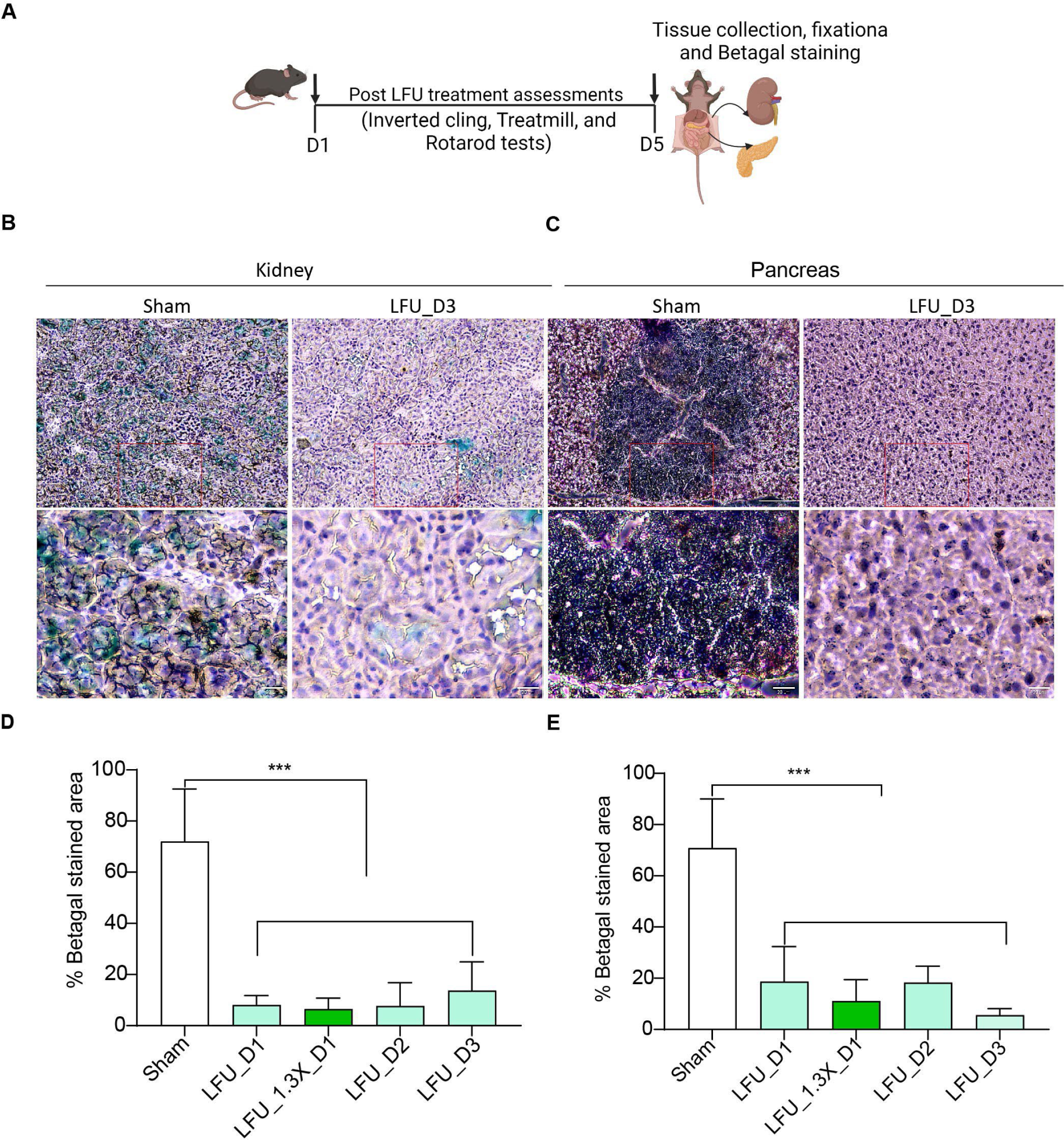
LFU decreases fraction of SA-β galactosidase staining cells in kidney and pancreas. (A) Schematic illustration of the treatment plan and tissue staining. Mice were treated 30 min every day with 1X power (LFU_D1) or 1.3X power, (LFU_1.3X_D1), every other day (LFU_D2) and every third day (LFU_ D3) day with 1X power for two weeks then two weeks break followed by a second two-week treatment. Performance of treated mice was assessed as pre and post assay (see Figure S18). Each study group contained five male and five female mice. After 4 weeks of treatment followed by assessment, mice were euthanized, and kidney and pancreas were collected for SA-β galactosidase staining. (B) SA-β-galactosidase-stained kidney sections of sham and LFU treated mice. (C) SA-β-galactosidase-stained pancreas sections of sham and LFU treated mice. Blue color indicates b-galactosidase activity, scale bar=150 µm and 20 µm. (D) Quantification of b-galactosidase staining of kidney sections in sham and LFU treated mice. (E) Quantification of b-galactosidase staining of pancreas sections in sham and LFU treated mice. Minimum ten mice per group were used otherwise mentioned. Results are plotted as mean ± S.D. Two tailed unpaired Student t-test was used to determine the statistical significance. P-value greater than 0.05 is represented by ns. Statistical significance was given in p-value. *P<0.05, **P<0.002, *** P<0.0001.

## Notes

### Summary of Updates

We have updated the new data into our previous manuscript.

